# Fitness defects due to cytosolic protein misfolding in *S. cerevisiae* can be alleviated by decreasing mitochondrial protein import capacity

**DOI:** 10.1101/2024.08.26.609637

**Authors:** Zainab Zaidi, Devi Prasanna Dash, Akanksha Sharma, Soumen Kundu, Sarika Bhatt, Shikha Rao, Kedar Padia, Manish Rai, Kausik Chakraborty

## Abstract

Protein misfolding affects cellular fitness. This can be caused due to the toxic aggregation of one species of protein or global protein misfolding events. Since the fitness defect arises due to the multi-modal effect of misfolding, there is no consensus mechanism to alleviate this fitness defect. Here, we used adaptive laboratory evolution of thermotolerance to identify pathways contributing to proteotoxic stress resistance in *S. cerevisiae*. Our results suggest a link between thermotolerance and proteotoxicity resistance, majorly routed through the loss of mitochondrial DNA. Loss of mitochondrial DNA decreased the association of mistargeted misfolded proteins on the mitochondrial surface and altered the cellular response to proteostasis to enhance protein quality control associated degradation. We show that a decrease in the abundance of import channels is sufficient to mimic the loss of mtDNA and increase cellular proteostasis. Thus, we uncover a cryptic interorganellar cooperation in combating proteotoxicity in yeast.

## INTRODUCTION

Most polypeptide chains must fold into their native state to function correctly. Although the information necessary for achieving the correct structure is encoded within the polypeptide itself, many proteins require assistance to attain this structure within a physiologically relevant timeframe while avoiding aggregation^1^. To facilitate proper folding, all cells and organisms have evolved a complex network of proteins known as chaperones^2,3^. These chaperones assist in the folding of proteins that require assistance. When proteins fail to reach their native form, the protein quality control network recognizes them and degrades or sequesters them into inert aggregates ^4,5^. Given this elaborate system, it is surprising that protein misfolding is often associated with dominant toxicity.

While recessive mutations may lead to fitness loss as proteins lose their function, mainly when a specific protein is vital for cellular health, accumulation of non-native forms of certain proteins can be toxic to cells^6–9^. This toxicity is not because the mutated protein lost its native function but because the mutant protein is toxic in the non-folded state^8,10^. Numerous studies over the past few decades have illuminated the toxicity mechanisms in different disease models, suggesting the smaller oligomeric aggregation intermediates to be more harmful than the final amyloid structures^11–13^. These toxic oligomers interact with many proteins, including essential chaperones^14,15^, reinforcing that these aggregates can sequester crucial fitness-defining proteins reducing cellular fitness^16,17^. Moreover, there is evidence that these oligomers can be toxic to membranes^18^ or interact with sub-cellular organelles, potentially disrupting their homeostasis and biogenesis^19–21^. Consequently, the underlying basis for this toxicity is complex, and the relative contribution of each in reducing overall fitness, is not known.

Since chaperones are sequestered by the toxic oligomeric intermediates, overexpressing certain chaperones should help mitigate toxicity. While there is an expanding body of literature focused on chaperones that mitigate these toxic effects, a common determinant that can alleviate proteotoxic stress has yet to be identified^22,23^. An extensive genetic screen conducted by Silva et al. revealed numerous genes that could reduce protein aggregation or the toxicity of misfolding in *C. elegans*^24^. However, it has proven challenging to establish a common theme that effectively counters misfolding-induced toxicity. Like other model organisms, budding yeast (*S. cerevisiae*) has been extensively utilized to identify genetic modulators of protein aggregation-associated toxicity. Most of these studies have employed toxic protein aggregates, such as PolyQ expansion models of Htt, alpha-synuclein, transthyretin, and A-beta, to uncover genetic modifiers^25,26^. While these findings are informative, it is equally critical to understand the common pathways that can mitigate misfolding-associated stress, even when the misfolding protein does not form amyloid aggregates. Without this knowledge, it is difficult to understand if a common strategy can be engineered to deal with dominant toxic misfolding events.

To address this need, we aimed to employ a different genetic approach to identify potential modifiers of proteotoxic stress. Commonly, genetic modifiers in yeast and *C. elegans* are identified using reverse genetics^24,25,27–29^. While this method is extremely effective, it often overlooks the complex interplay of different modifiers. Therefore, we pursued a forward genetics approach, using growth at high temperatures as the selection criterion. Adaptive laboratory evolution (ALE) strategies have been instrumental in elucidating the molecular details of evolution and the paths taken during short-term adaptations^30–34^. With advancements in genomic tools, it is now feasible to utilize ALE to select for desirable traits and identify molecular modulators through high-throughput sequencing. We hypothesized that conducting ALE at elevated temperatures might favor strains capable of coping with the toxic effects of protein misfolding and aggregation. Although high-temperature growth is associated with various stresses, protein misfolding and aggregation pose significant challenges that cells must overcome to survive^35^. By conducting multiple parallel evolution experiments, we aimed to identify the most successful pathways and pinpoint modifiers with high confidence, especially when replicated across multiple ALE trials.

We discovered the loss of mitochondrial DNA (mtDNA) as a predominant and consistent event during ALE at high temperatures. We confirmed this loss was linked to altered proteostasis and an enhanced ability to manage misfolding stress. We then explored whether this information could be used to identify (1) the primary contributor to fitness defects during misfolding stress and (2) a pathway that could alleviate this stress. We found that cytosolic misfolded proteins associate with mitochondria for degradation through the canonical mitoTAD^36^ or MAD^37^ pathways, which become toxic to mitochondria during excessive misfolding stress. Additionally, we observed that preventing mitochondrial import can enhance the degradation of misfolded proteins by increasing proteasomal flux without affecting the concentration of proteasomes themselves. Our findings highlight mitochondrial import quality control pathways as critical components in maintaining cytosolic and overall cellular proteostasis. While these pathways serve as guardians, they can become vulnerabilities in the face of high misfolding loads. We propose that the import channels have evolved to fulfill multiple roles within the cell, including a crucial function in overall proteostasis.

## RESULTS

### Laboratory evolution of thermotolerant *S. cerevisiae*

We used a well-established laboratory evolution methodology to evolve the thermotolerant yeast cells through serial dilution. The details of the evolution experiment are published elsewhere^38^. To uncover the most favored routes for evolving thermotolerance, we performed 29 parallel evolution experiments at 40^0^C, with the haploid *Saccharomyces cerevisiae* parent strain BY4741 (Figure 1A). After 600 generations of evolution, we randomly isolated a single colony from each evolved population and examined their growth rates at 40°C and 30°C. The growth rates at 40^0^C were higher than the parental strain’s, confirming that the strains evolved thermotolerance (Figure 1B, S1A). To further assess the thermotolerance of these strains, we monitored their competitive fitness at 40°C. For this, we designed a fluorescence-based competitive fitness assay. The detailed design of the experiment is shown in (Figure 1C). Briefly, we mixed a GFP-tagged parent strain (BY4741 with a genome-integrated GFP) with an untagged evolved strain in a 1:1 cellular ratio and grew the mixed culture at 30°C or 40°C. The ratio of the number of non-fluorescent-cells (evolved) to fluorescent cells (parental) at different times of growth, provided a quantitative measure of the fitness advantage of the evolved strains. To identify a GFP-tagged BY4741 that has the same fitness at 40^0^C as that of un-tagged parental BY4741, we screened for multiple GFP-tagged proteins available in Yeast GFP Clone Collection^39^ and found C-terminal GFP tagged TDH2 suitable for the assay. It had the same fitness as BY4741 at 40^0^C and exhibited a homogenously fluorescent population at that temperature (Figure S1B). For all fitness experiments, BY4741 with TDH2-GFP was used as a proxy for parental BY4741. Competition between BY4741(TDH2-GFP) and BY4741 was always included as a control (Figure S1B, S1C). The evolved strains displayed higher competitive fitness at 40°C than the parental strain (Figure 1D). Further details confirming the evolution of thermotolerance and fixation along with control experiments are reported in detail in an accompanying manuscript^40^. Thus, we generated strains that were fitter than the parental strain at 40^0^C providing us with a handle to ask if thermotolerance is linked with the evolution of better proteostasis.

**Figure 1.**
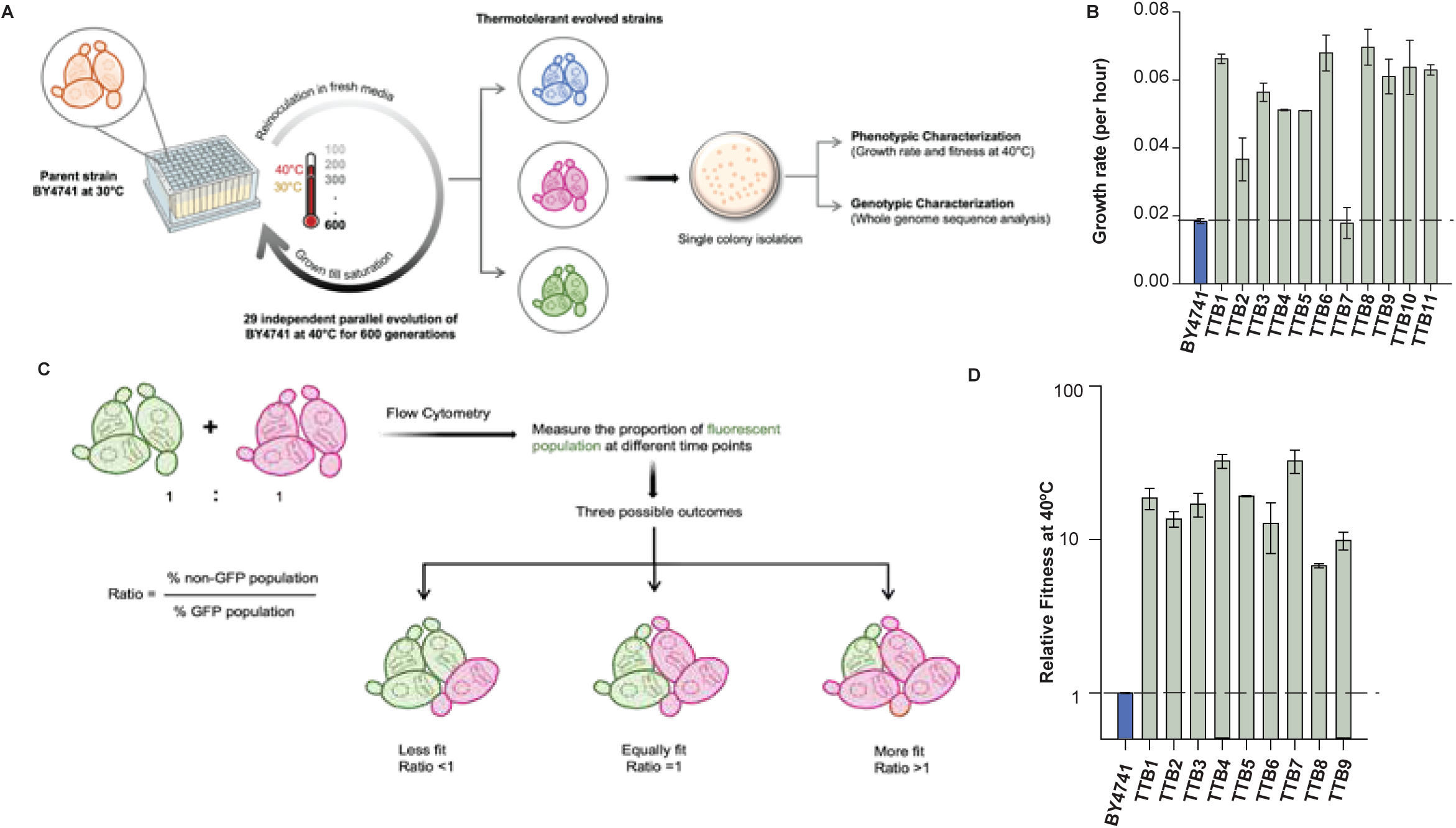
Yeast cells acquired thermotolerance in 600 generations. A) Schematic describing the experimental outline of adaptive laboratory evolution (ALE). ALE experiment starts with the parental strain BY4741, subjected to high temperature (40°C) in YPD. The cells were grown till saturation and re-inoculated in fresh media. This was followed for ∼600 generations, after which single colonies were isolated from each independent pool of evolved strains and were characterized phenotypically and genotypically. B) Growth rate of resultant evolved strains at 40°C in YPD medium. TTB1, TTB2, TTB3, TTB4, TTB5, TTB6, TTB7, TTB8, TTB9, TTB10, TTB11 correspond to evolved thermotolerant strains and BY4741 is the parent strain. C) Experimental design of competitive fitness assay. Briefly, the fluorescently tagged (TDH2-GFP) parental strain is mixed with untagged evolved strain in equal cellular ratio. Following their growth at the specified conditions, the proportion of each population was measured using flow cytometry and the relative fitness was assessed using the ratio of both the populations. D) Fitness of evolved strains at 40°C confirming the acquisition of thermotolerance. The data is normalized with respect to the parent strain BY4741 and the fitness of respective strains at 30°C.

### The evolution of yeast at high temperatures can take two major routes

We sequenced the genomes of all the evolved strains to identify common genetic routes to evolving thermotolerance in BY4741. We identified point mutations in the genes, *lrg*1^41^ and *mrn*1^42^, whose loss of function is known to increase thermotolerance. We also found disomy of chromosome III in multiple strains which recapitulated earlier observations in diploid *S. cerevisiae* (Figure 2A)^32^. Thus, we were able to sample the known routes to achieve thermotolerance in *S. cerevisiae*. Interestingly, loss of mtDNA reads was the most predominant feature in the evolved strains; ∼75% percentage of the evolved strains showed this phenotype (Figure 2A). The two apparent solutions to thermotolerance, loss of mtDNA and disomy of Chromosome III, were exclusive solutions with no strains having both the features (Figure 2A; right panel). While the average depth of mtDNA reads was negligible, we confirmed the loss using quantitative PCR against a mitochondrially encoded gene, *COX-3* (Figure 2B). To differentiate between a segmental deletion in mtDNA and a complete loss, we analyzed the read depth throughout the mitochondrial genome and found the read depths to be uniformly low or absent in the evolved strains lacking mtDNA (Figure 2C and Figure S2). Furthermore, these strains grew poorly on non-fermentative carbon sources, an indication of the non-respiratory phenotype associated with the loss of mtDNA (Figure 2D). This was unique to strains that lost mtDNA and not the ones with the disomy (Figure 2D). Thus, the evolution of thermotolerance adopted two mutually exclusive routes: ChrIII disomy and loss of mtDNA.

**Figure 2.**
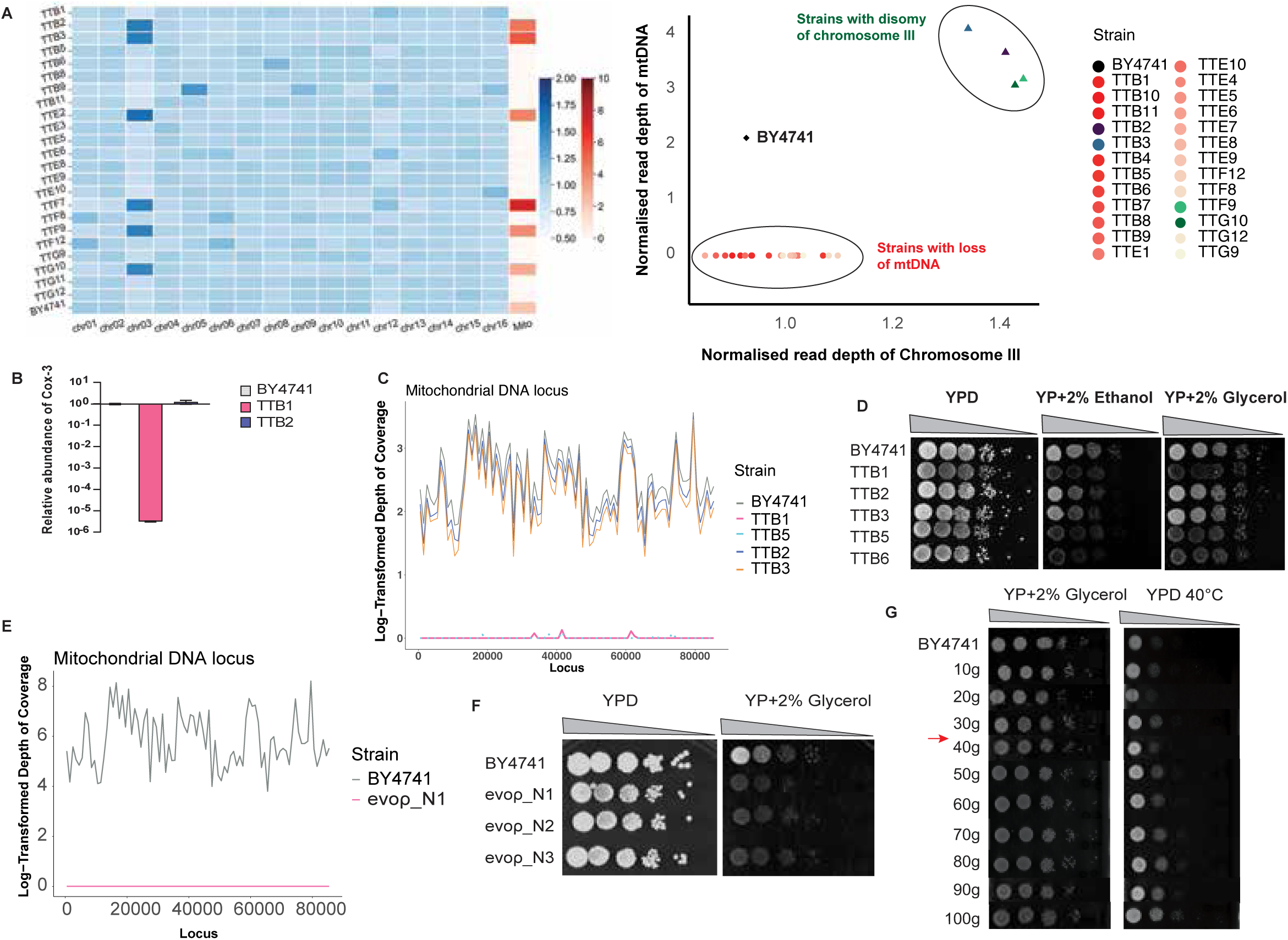
Evolution of thermotolerance in yeast results in two distinct outcomes. A) Heatmap representing the chromosome-wise depth of coverage in evolved thermotolerant strains. Blue scale indicates the nuclear genome depth of coverage and red scale represents the mitochondrial genome depth (left panel). Graph depicting strain-wise correlation between observed disomy of chromosome III and loss of mtDNA (right panel). B) Abundance of mitochondrial gene COX-3 in representative thermoevolved strains from both categories. Ct values are normalized wrt parent strain BY4741. Mean is plotted with standard deviation as error bar from three biological replicates. p-value< 0.05. C) Depth of coverage at each locus of the mitochondrial genome in thermotolerant strains. Pink and Cyan line represents TTB1 and TTB5, strains with loss of mtDNA, depicting complete loss of mtDNA. Value +1 was added to the mean depth of chromosomes and log2 transformed. D) Spot assay showing impaired respiration in TTB1, TTB5 and TTB6 relative to parent strain BY4741 and other thermotolerant strains with disomy of chromosome III (TTB2, TTB3). Log-phase cultures were serially diluted and spotted on fermentative (glucose) and non-fermentative carbon sources (glycerol and ethanol) to assess respiratory function of the cells. E) Depth of coverage in evoρ^0^ N1 strain throughout the mitochondrial genome, depicting complete loss of mtDNA in independent evolution experiment. Value +1 was added to the mean depth of chromosomes and log2 transformed. F) Spot assay on non-fermentative carbon source (glycerol) to assess respiration impairment in evoρ^0^ strains. (n=3) G) Spot assay showing the association between loss of mtDNA and acquisition of thermotolerance in intermediate generations.

### Loss of mtDNA is a reproducible outcome of evolution

To check if the loss of mtDNA is a reproducible event during the evolution of thermotolerance, we repeated the laboratory evolution of BY4741 at 40^0^C for 100 generations and stored cultures every tenth generation in between. Following the evolution, we ascertained that the strains acquired thermotolerance in a span of 100 generations (Figure S3A and S3B). Functional and genetic characterization of the freshly evolved strains revealed that the cells lost mtDNA, hereafter referred to as evoρ^0^ (evolved thermotolerant strains without mtDNA) strains (Figure 2E and 2F, Figure S3C and S3D). This confirms that the loss of mtDNA is a reproducible outcome of evolution at high temperatures. Additionally, we find a strong association between thermotolerance and loss of mtDNA by monitoring these at the intermediate generations (Figure 2G and S3E).

We also subjected two genetically distinct strains of *Saccharomyces cerevisiae* (W303, RM11-1a) to high-temperature ALE to investigate whether loss of mtDNA is a general outcome. While the evolved W303 strains lost mtDNA, the evolved RM11-1a strains retained it (Figure S3F). Interestingly, BY4741 and W303 are known to have a T661A substitution in MIP1 (mitochondrial DNA polymerase) that makes the protein thermolabile^43^. This leads to mtDNA loss at high temperatures due to exacerbated mutation accumulation in the mitochondrial genome. RM11-1a lacks this substitution, resulting in stable mtDNA and fewer petites (Figure S3F; right panel). This data helps us in explaining how mtDNA is lost at higher temperatures in BY4741. However, even with the same mutation on MIP1, the loss of mtDNA in W303 was still much less frequent than BY4741 (1 out of 3 replicates in W303 vs 3 out of 3 replicates in BY4741), indicating a selective advantage of losing this in the background of BY4741. Therefore, we asked if the loss of mtDNA provides this advantage by reducing proteotoxicity associated with elevated temperatures.

### Loss of mtDNA makes cells resilient to proteotoxic stress

Next, we asked if the evoρ^0^ strains have better proteostasis. To assess the proteostasis capacity of evoρ^0^ strains, we used L-Azetidine-2 Carboxylic Acid (AZC). AZC is a proline analog that is efficiently incorporated in polypeptide chains altering protein conformations^44,45^. This incorporation led to misfolding as indicated by the upregulation of canonical heat shock response in BY cells treated with AZC (Figure S4A and S4B). Additionally, we saw an increase in protein ubiquitination (Figure S4C) and aggregate formation (Figure S4D) upon treating the cells with AZC, indicating that this molecule induces global misfolding stress. evoρ^0^ strains grow better than the parental strain (BY4741) in the presence of AZC (Figure 3A). Similarly, evoρ^0^ strains also showed significantly higher fitness in AZC than BY4741, suggesting that evoρ^0^ strains have an enhanced ability to cope with global proteotoxic stress induced by AZC (Figure 3B). Since evoρ^0^ strains from both evolution experiments showed higher fitness under global proteotoxic stress conditions (Figure 3B and Figure S4E), this implied a correlation between the loss of mtDNA and the emergence of better proteostasis capacity. We hypothesized that the emergence of AZC tolerance should coincide with the loss of mtDNA during evolution. During ALE, mtDNA loss commenced as early as the 20th generation and became more homogenous by the 40th generation (Figure S4F and 2G). As anticipated, AZC tolerance emerged as the cells lost mtDNA, underlining the correlation between the two (Figure S4G). Single colonies of 20^th^, 30^th^ and 40^th^ generations showed a similar correlation between mtDNA loss and AZC tolerance (Figure 3C). In the 20^th^ generation, a mixed population of cells with and without mtDNA was observed (20ρ^+^ and 20ρ^0^, respectively). Even at the same stage of the evolution experiment, the 20ρ^0^ strain had higher AZC tolerance than the 20ρ^+^ strain (Figure 3D and 3E). This data suggests a strong correlation between mtDNA loss and better proteostasis capacity.

**Figure 3.**
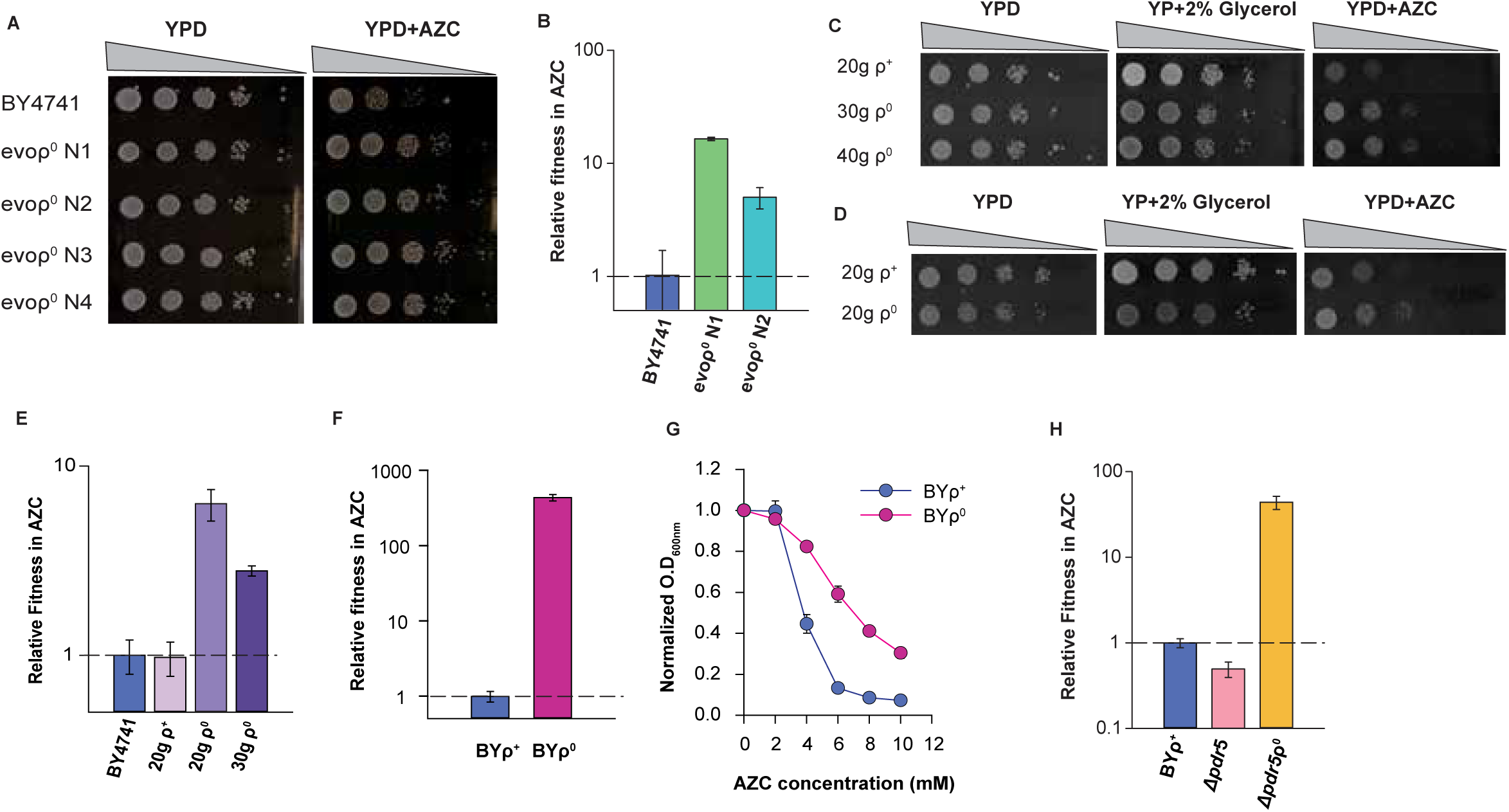
Reconstitution of ρ^0^ phenotype alone is sufficient to alter the proteostasis capacity of the cell. A) Spot assay showing the growth differences between the parent (BY4741) and evolved strains (evoρ^0^ N1, evoρ^0^ N2 and evoρ^0^ N3) on AZC-containing media. Log phase cultures were serially diluted and spotted on YPD plates with and without 5mM AZC at 30°C. (n= 4) B) Relative fitness of evoρ^0^ strains in 4mM AZC in YPD at 30°C after 48 hours. Normalization is done wrt parent strain (BY4741). Error bars represent standard deviation from three replicates. p-value< 0.05, calculated using Student’s t-test. C) Spot assay showing the concomitant loss of mtDNA and gain of AZC tolerance in the intermediate generations of evolved strains (20, 30 and 40 generations). The presence (ρ^+^) or absence (ρ^0^) of mtDNA is indicated (n=3). D) Single colonies from the 20th generations showing the correlation between mtDNA loss and AZC tolerance. (n=3) E) Relative fitness of evolved strains (with or without mtDNA loss) in 4mM AZC at 30°C, normalized wrt BY4741. n=3, p-value< 0.05 calculated using Student’s t-test. F) Relative fitness of BYρ^0^ in 4mM AZC containing YPD at 30°C after 48 hours. The fitness of the BYρ^0^ is normalized wrt BYρ^+^ (BY4741). Error bars indicate standard deviation from three replicates. Statistical significance calculated using Student’s t-test, p-value< 0.05. G) MIC assay of BYρ^0^ and BYρ^+^ in increasing concentration of AZC in YPD at 30°C. H) Relative fitness of deletion strain (Δ*pdr5*) and its ρ^0^ counterpart (Δ*pdr5*ρ^0^) relative to BY4741 in AZC. n=3, error bars are plotted by calculating standard deviation from three biological replicates with p-value <0.05.

In addition to the absence of mtDNA, the evoρ^0^ strains harbor mutations in their nuclear genome. To deconvolute the role of ρ^0^ phenotype from co-occurring mutations, we generated ρ^0^ on the parental strain background BY4741 (BYρ^+^) , referred to as BYρ^0^. This approach bypasses evolution and negates the role of nuclear mutations that accumulated during ALE, providing us with a cleaner background to study the effect of mtDNA loss on proteostasis. As expected, BYρ^0^ exhibited more than a 10-fold increase in fitness over BYρ^+^ in AZC (Figure 3F, Figure S4H). Additionally, BYρ^0^ also showed more tolerance to AZC in MIC assay where the strains (BYρ^+^ and BYρ^0^) were grown in increasing concentrations of AZC (Fig 3G). Trivially, BYρ^0^ strains are known to increase the efflux of toxins by upregulating PDR5^46^. Even in the Δ*pdr5* background, loss of mtDNA increased resistance to AZC (Figure 3H), indicating that AZC resistance of BYρ^0^ is not due to enhanced AZC efflux. Collectively, these findings demonstrate that the loss of mtDNA alone is sufficient to alter the proteostasis capacity of the cell.

### **ρ** strains have enhanced degradation capacity through proteasome

To understand why cells devoid of mtDNA are resilient to proteotoxicity, we checked if loss of mitochondrial translation confers AZC resistance. Since mtDNA encodes genes that are translated in mitochondria, AZC may be incorporated in the mtDNA-encoded proteins. If these are especially toxic for the cell, removing mtDNA would relieve growth defects arising from AZC treatment. To test this, we specifically inhibited mitochondrial translation in BYρ^+^ and checked if it conferred AZC resistance. No fitness advantage in AZC was seen upon inhibition of mitochondrial translation, negating its role in enhanced proteostasis capacity of BYρ^0^ (Figure S5A).

To decipher the mechanism, we systematically probed the different arms of the proteostasis network (PN): protein folding, translation, degradation, and aggregation. First, we checked if the loss of mtDNA increased the folding capacity of BYρ^0^ over BYρ^+^. To investigate this, we utilized a cytosolic folding reporter protein, TS22^47^. This protein is a destabilized version of Nourseothricin acetyl-transferase (Nat-R) that confers Nourseothricin (CloNAT)-resistance to *S. cerevisiae*. TS22 exhibits lower activity in BYρ^0^ strains than in BYρ^+^without altering the activity of WT Nat-R (Figure 4A). Similarly, even between the thermotolerant strains, TS22 had lower activity in 20ρ^0^ than in 20ρ^+^ (Figure S5B). This was not specific for TS22, as a less severe mutant of Nat-R, TS15^47^, also showed lower activity in BYρ^0^ than in BYρ^+^ (Figure S5C). This suggested that the loss of mtDNA did not increase the folding capacity of BYρ^0^. On the contrary, a decrease in the activity of the mutants suggested a decrease in protein translation or an increase in protein degradation capacity. WT Nat-R did not show a difference in activity (Figure 4A) indicating that expression from the promoter did not change transcriptionally or translationally in the ρ^0^ strains. To quantitatively rule out any difference in translation as the underlying reason for AZC tolerance, we measured single-cell fluorescence of mCherry fused to WT Nat-R. BYρ^0^ did not show a lower expression level of Nat-R-mCherry than BYρ^+^ (Figure 4B), ruling out altered expression or translation as the reason for decrease in activity of Nat-R mutants in BYρ^0^ strains. We also confirmed similar translation capacity in BYρ^+^ and BYρ^0^ by measuring the expression levels of GFP driven by a different promoter (Figure 4B). Thus, reduced translation was not the reason for the enhanced ability of BYρ^0^ to tolerate proteotoxic stress.

**Figure 4.**
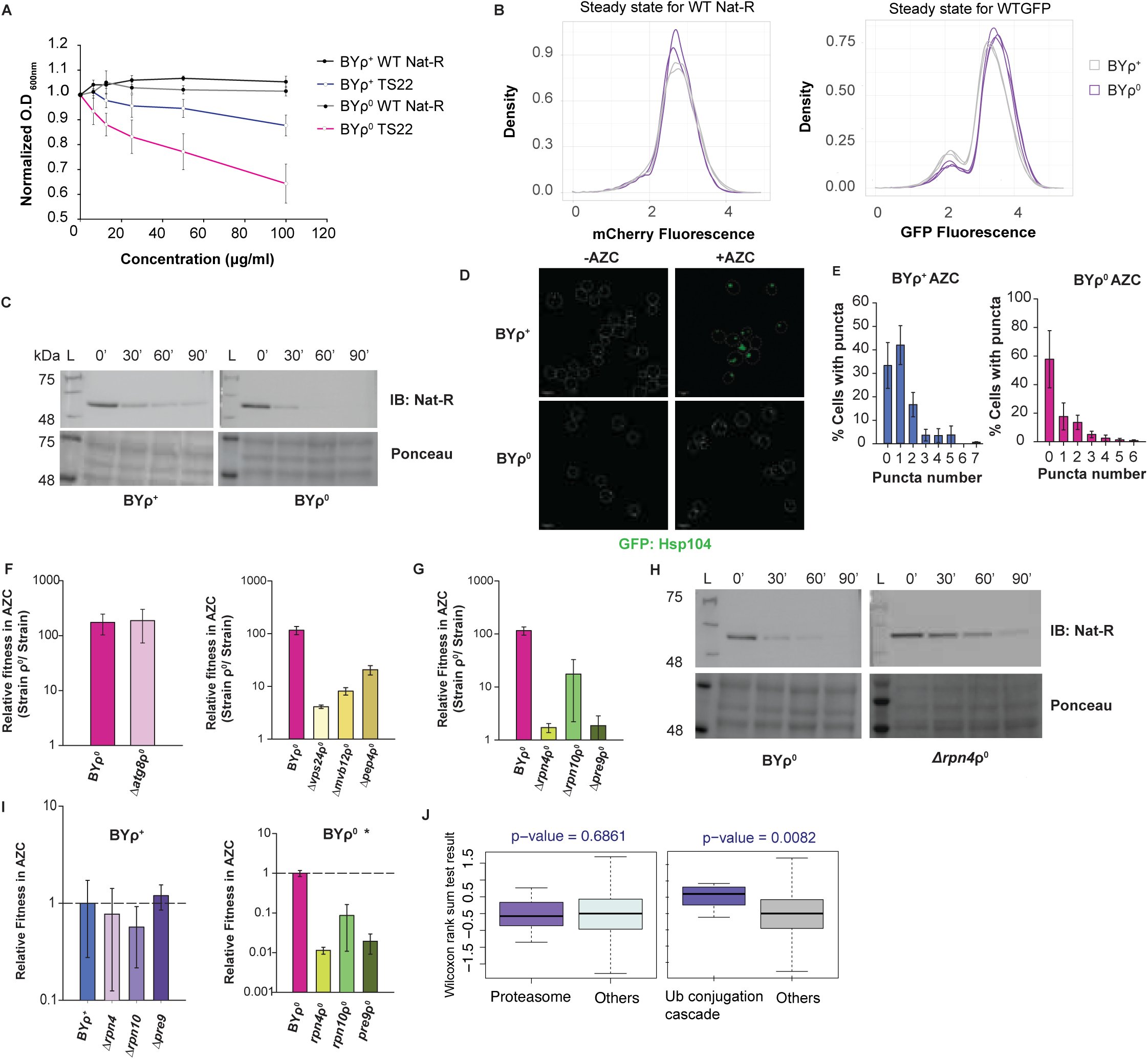
BYρ^0^ have enhanced degradation capacity potentially mediated through proteasome. A) Growth assay of WT Nat-R and TS22 in BYρ^+^ and BYρ^0^ with increasing concentrations of nourseothricin (Clonat) at 30°C to assay for the functionality of the proteins. (n=3) B) Histogram depicting the expression of WT Nat-R (left panel) and WTGFP (right panel) in BYρ^+^ and BYρ^0^. Nat-R and WT GFP are driven by TEF and GPD promoters respectively. Fluorescence of mCherry is used as a proxy for expression of WT Nat-R as WT Nat-R is fused with mCherry. C) Cycloheximide chase assay of TS22 in BYρ^+^ and BYρ^0^ to monitor the rate of degradation of the misfolded protein at 30°C. Ponceau is used for normalization of total protein (loading control) n=4. D) Aggregation in BYρ^+^ and BYρ^0^ upon AZC treatment at 30°C. GFP signal corresponds to HSP104. E) Quantification of D) F) Relative fitness of ρ^0^ deletion strains from autophagy pathway (left panel) and vacuolar pathway (right panel) in AZC. Data is normalized relative to respective ρ^+^ deletion strain. Error bars represent standard deviation from three biological replicates. p-value; ns for autophagy. <0.05 for vacuolar G) Relative fitness of ρ^0^ deletion strains of proteasome pathway in AZC. Normalization is done wrt respective ρ^+^ deletion strain. Standard deviation is plotted as error bars and p-value is calculated using Student’s t-test. p-value<0.05, n=3 H) The stability of TS22 in BYρ^0^ and Δ*rpn4*ρ^0^ at 30°C. Cells were grown till lo-phase and treated with 40μg/mL cycloheximide to arrest translation. Different time points were collected and protein lysate was loaded. Ponceau is used as loading control. I) Relative fitness of proteasomal gene deletion strains (left panel) and ρ^0^ deletion strains in AZC. Normalization is done with BYρ^+^ and BYρ^0^ respectively. Mean is plotted with standard deviation as error bars. Significance is calculated using Student’s t-test. p-value; ns for left graph and <0.05 for right graph. * denotes same data as G), represented differently. J) Statistical significance of alterations in proteasomal subunits (left panel) and ubiquitin conjugation cascade (right panel) in BYρ^0^ with AZC treatment, compared to BYρ^+^ with AZC treatment. Significance was calculated using Wilcoxon rank-sum test, with p-value indicated on the graph.

Since the activity of Nat-R mutants was low in BYρ^0^, we examined if the Nat-R mutants were degraded faster in BYρ^0^ than in BYρ^+^. We performed Cycloheximide (CHX) chase assay for TS22 in both BY and BYρ^0^. Interestingly, TS22 turned over faster in BYρ^0^ than in BYρ^+^ while WT Nat-R showed no difference (Figure 4C, S5D). A similar result was obtained for TS15 (Figure S5E), although the difference was not as pronounced as TS22 (TS15 being a milder mutant). This demonstrated that BYρ^0^ had an enhanced capacity to degrade misfolding protein proteins while sparing folded ones, a hallmark of improved protein quality control. To ensure that TS22 doesn’t depend upon a special route to be turned over and is indeed turned over through canonical protein quality control pathways, we monitored the turnover of TS22 while treating the cells with AZC (Figure S5F). There was a drastic reduction in TS22 turnover indicating that it takes the protein quality control associated route to degradation of endogenous proteins; when there is excess misfolding of endogenous proteins, they compete out the existing channels of degradation to decrease the turnover of TS22. To confirm that the enhanced clearance rate in BYρ^0^ is not confined to NAT mutants, we followed the fate of endogenous misfolded proteins during AZC treatment; these are typically marked for degradation by ubiquitination. Ubiquitinated proteins turned over faster in BYρ^0^ than in BYρ^+^ (Fig S5G) confirming that even endogenous misfolded proteins are degraded more efficiently in BYρ^0^. Thus, BYρ^0^’s ability to tolerate proteotoxicity may be linked to its increased ability to clear out misfolded proteins.

Additionally, a strain may also be resilient to proteotoxicity if it can better package the misfolded proteins into inert protein aggregates^48^. To visualize aggregates in-vivo, we utilized GFP-tagged HSP104. Interestingly, aggregation status differs drastically between BYρ^+^ and BYρ^0^ in response to AZC (Figure 4D), while there was no appreciable difference between the untreated cells. The aggregates in the BYρ^0^ strain were less abundant and smaller in size as compared to BYρ^+^ (Figure 4E). This demonstrates that BYρ^0^ does not use efficient sequestration of misfolded proteins to counter proteotoxicity; on the contrary, this resonates with the earlier observation that BYρ^0^ can efficiently clear off misfolded proteins, leaving little to form aggregates. These findings also indicate that the increased degradation capacity of BYρ^0^ is not limited to the Nat-R mutant protein alone but extends to the context of global proteotoxic stress, where misfolded proteins are cleared more rapidly. Taken together, only the degradation arm of the proteostasis network is enhanced in BYρ^0^ with no appreciable changes in protein translation, folding, or aggregation branches.

To decipher the degradation pathway responsible for the faster turnover of misfolded proteins in BYρ^0^, we deleted the key components of canonical degradation pathways in BYρ^0^ and examined the fitness of the resulting strains in the face of global proteotoxic stress. Deletion of a pathway that contributes to the enhanced degradation in the BYρ^0^ strain, should reduce the fitness advantage of ρ^0^ in AZC. While the deletion of regulatory genes of vacuolar degradation (Δ*vps24,* Δ*mvb12,* Δ*pep4*) and autophagy pathway (Δ*atg8*) contributed little to the fitness advantage of BYρ^0^ in AZC (Figure 4F), the deletion of proteasomal genes (Δ*rpn4,* Δ*rpn10,* Δ*pre9*) produced the most striking fitness defect (Figure 4G). This underlined the involvement of the proteasomal degradation pathway in conferring AZC tolerance to BYρ^0^. In agreement with this, we found the degradation of TS22 Nat-R to be slower in the Δ*rpn4*ρ^0^ strain than in BYρ^0^ (Figure 4H). Although BYρ^0^ depended on the proteasomal genes for AZC tolerance, BYρ^+^ showed no such dependence on the proteasomal components (Figure 4I). Consequently, the fitness of BYρ^+^ and BYρ^0^ in AZC becomes comparable in the background of these deletions. This suggested that BYρ^0^ depends more on the proteasomal components than BYρ^+^ for efficiently tolerating global misfolding events.

To check if this is due to a higher abundance of proteasomal components in BYρ^0^ than in BYρ^+^, we performed label-free comparative proteomics of these two strains in the presence and absence of AZC. Notably, the abundances of the subunits were not different between BYρ^+^ and BYρ^0^ in the presence of AZC (Figure 4J;left panel). Thus, the enhanced ability of BYρ^0^ to degrade proteins during AZC stress was not a result of an increase in the abundance of the proteasome. However, in AZC, the levels of E1/E2/E3 enzymes were significantly upregulated as a gene set, in BYρ^0^ than in BYρ^+^ (Figure 4J; right panel). Thus the response of BYρ^0^ to AZC may ensure a more efficient misfolded protein turnover than in BYρ^+^ even in the presence of a similar concentration of proteasomes.

### Misfolded proteins associate with mitochondria in BY

Other non-canonical pathways involving mitochondria may be involved in the degradation of cytosolic misfolded proteins. Two pieces of evidence, other than the loss of mtDNA in the evolved strains hinted towards the role of mitochondria. First, we found mitochondrial fragmentation to be significantly higher during AZC stress (Figure S6A), indicating mitochondrial stress. Second, although the deletion of *rpn4*, the master regulator of the proteasome, did not increase sensitivity of BYρ^+^ towards AZC, the deletion of *doa1* (a protein involved in mitochondria-associated degradation) increased the sensitivity (Figure S6B; S6C).

Mitochondrial degradation of cytosolic misfolded proteins was recently reported to play an essential role during proteotoxic stress and was named Mitochondria As Guardians In Cytosol (MAGIC)^49^. We checked if BYρ^0^ depended on this pathway to avert proteotoxicity. MAGIC depends on the H^+^ potential difference across the mitochondrial inner membrane that is dissipated with CCCP that is otherwise inert towards protein degradation by proteasome or proteases. If BYρ^0^ depended more on this pathway than BYρ^+^, CCCP treatment should decrease the degradation rate of TS22 in BYρ^0^ and should make it comparable to BYρ^+^. Unexpectedly, the opposite was observed (Figure 5A); there was a stark decrease in the degradation of TS22 in BYρ^+^ while the degradation rate in BYρ^0^ remained relatively unchanged. Inhibition of respiration with oligomycin did not show such a stark difference (Figure S6D) highlighting the importance of proton gradient over the electron transport chain for the degradation of TS22. Contrary to our expectations, this suggested that mitochondrial potential may play a role in misfolded protein degradation in BYρ^+^ but not in BYρ^0^. Although this observation countered our working hypothesis, we posited that this difference may play a role in dictating the proteotoxicity tolerance of BYρ^0^. A recent report suggested that MAGIC during metabolic stress could decrease cellular fitness^50^. In line with this, we speculated that lower MAGIC in BYρ^0^ than in BYρ^+^ may contribute to its higher fitness in AZC. We reasoned that misfolded protein association with mitochondria may be toxic during global misfolding stress thereby making BYρ^+^ more sensitive to this proteotoxic insult than BYρ^0^.

**Figure 5.**
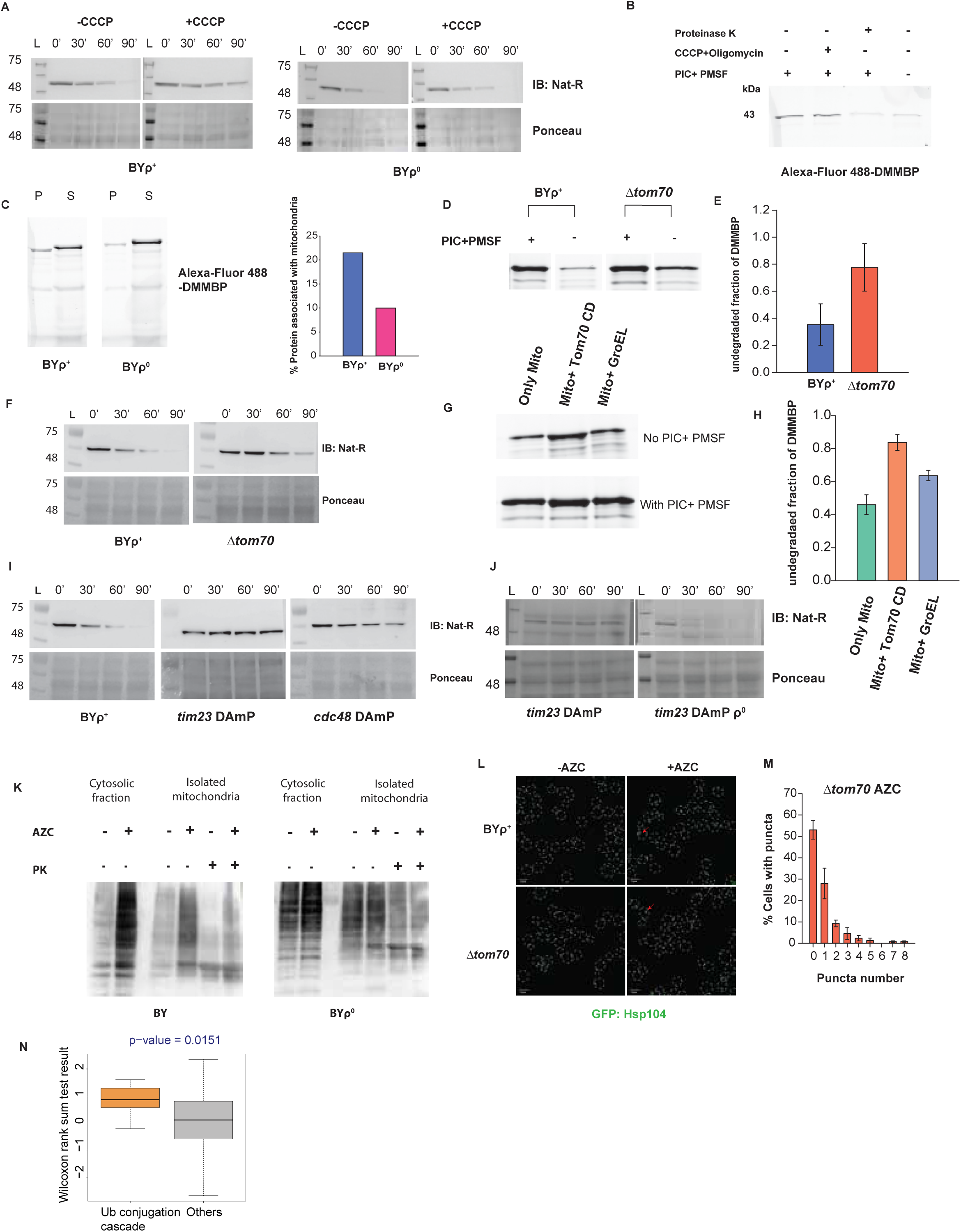
Misfolded protein associates with mitochondria. A) Cycloheximide chase assay of TS22 in BYρ^+^ (left panel) and BYρ^0^ (right panel) in the presence of 50uM CCCP. Ponceau is used for normalization of total protein. B) in-vitro association of unfolded DMMBP with isolated mitochondria of BYρ^+^. DMMBP labelled with Alexa-Fluor 488 is utilized for the assay. Image shows the pellet fraction. C) in-vitro association of unfolded DMMBP with isolated mitochondria of BYρ^+^ and BYρ^0^. P corresponds to the pellet fraction and S corresponds to the supernatant fraction (left panel). Quantification of the same is shown in the right panel. D) In-vitro assay demonstrating the reduced degradation of unfolded DMMBP protein in isolated mitochondria of Δ*tom70* compared to BYρ^+^. E) Quantification of D). The undegraded fraction of DMMBP was calculated as the ratio of protein amount remaining without treatment (no PIC and PMSF) to the amount with treatment of PIC and PMSF. (n=4) F) Cycloheximide chase assay of TS22 in Δ*tom70* demonstrating the slower degradation of the protein as compared to BYρ^+^. Ponceau is used for normalization of total protein. G) In-vitro assay demonstrating the reduced degradation of unfolded DMMBP protein in isolated mitochondria of BYρ^+^ in the presence of cytosolic domain of Tom70 CD and GroEL respectively. H) Quantification of F). The undegraded fraction of DMMBP is calculated as the ratio of protein amount remaining without treatment (no PIC and PMSF) to the amount with treatment of PIC and PMSF. Bar plot depicts the average ratio of three biological replicates where error bars indicate the standard deviation I) Cycloheximide chase assay of TS22 in strains with downregulation of *tim23* and *cdc48*, demonstrating the slower degradation of the protein as compared to BYρ^+^. Ponceau is used for normalization of total protein. J) Cycloheximide chase assay of TS22 in *tim23* DAmP and *tim23* DAmP ρ^0^. Ponceau is used for normalization of total protein. K) Abundance of ubiquitinated proteins on the surface of isolated mitochondria from BYρ^+^ (left panel) and BYρ^0^ (right panel) and their respective cytosolic fractions. PK stands for Proteinase K. L) Aggregation in and BYρ^+^ and Δ*tom70* upon AZC treatment at 30°C. GFP signal corresponds to HSP104. M) Quantification of L) N) Statistical significance of alterations in ubiquitin conjugation cascade in Δ*tom70* with AZC treatment, compared to BYρ^+^ with AZC treatment. Significance was calculated using Wilcoxon rank-sum test, with p-value indicated on the graph.

For MAGIC, cytosolic misfolded proteins should be able to associate with mitochondria and enter mitochondrial compartments without a canonical mitochondrial import signal. To test this, we used purified mitochondria and checked if the non-native form of a slow-folding mutant of Maltose-Binding Protein (DM-MBP)^51^ was associated with mitochondria. This protein does not have a mitochondrial targeting signal. TS22 was not used for this assay as it was not amenable to in vitro folding studies. We started refolding DM-MBP in the presence of purified mitochondria by diluting out urea from denatured DM-MBP. DM-MBP was found associated with mitochondrial pellet (Figure 5B). It was found to be degraded in the absence of non-specific protease inhibitor, indicating that the binding is associated with turnover (Figure 5B). Notably, very little protein enters mitochondria; most of it stays on the surface of mitochondria and is accessible to proteinase K (Figure 5B). This suggested that the association does not lead to complete protein translocation inside mitochondria as required in MAGIC, but a transient association that nonetheless leads to protein turnover. The partitioning of DM-MBP to the mitochondrial pellet was markedly reduced in mitochondria isolated from BYρ^0^ (Figure 5C, S6E), supporting the previous evidence that mitochondria-associated degradation of TS22 was lower in BYρ^0^ (Figure 5A). Thus, we find a non-canonical mitochondria-associated degradation pathway in BYρ^+^ that is suppressed in BYρ^0^.

The receptors of unfolded protein on the surface of mitochondria (Tom70) could be essential for mitochondria-assisted degradation^52,49^. Indeed, when *tom70* was deleted, there was less degradation in vitro (Figure 5D, 5E, S6F), which was commensurate with decreased degradation of misfolded protein in vivo (Figure 5F). The specificity was further validated when the cytosolic domain of *tom70*, when added in vitro along with purified mitochondria from BYρ^+^ cells, was able to decrease the degradation of DM-MBP (Figure 5G, 5H, S6G). This was specific for the cytosolic domain of *tom70*, as GroEL, which can also bind to unfolded DM-MBP in an extended state, did not prevent its degradation as much as the *tom70* cytosolic domain(Figure 5G, 5H). Thus, the cytosolic domain of *tom70* and its anchorage on the mitochondrial membrane was critical in assisting the turnover of misfolded proteins on the mitochondrial surface. Interestingly, this was true for Tim23 downregulation (an import channel of the inner mitochondrial membrane) (Figure 5I) and partially with Cdc48 downregulation (Figure 5I), suggesting that the import channel and the extractase may play a role in clearing misfolded proteins associated with mitochondria. Significantly, in the background of BYρ^0^, depletion of Tim23 does not decrease the turnover of TS22 (Figure 5J), suggesting that BYρ^+^ was more dependent on mitochondria-assisted degradation than BYρ^0^. To check if this is true in chronic global misfolding stress, we monitored ubiquitinated proteins in the cytosolic and mitochondrial fraction of BYρ^+^ and BYρ^0^. Cytosolic and mitochondrially associated ubiquitination increased in BYρ^+^ upon AZC treatment, while only cytosolic ubiquitination increased in BYρ^0^ without a marked increase in the mitochondrial fraction (Figure 5K). The ubiquitinated proteins were on the mitochondrial surface and accessible to proteinase K (Figure 5K). Thus the import channels or receptors may be used to recruit misfolded proteins on the mitochondrial surface during AZC stress. This suggests that mitochondria-assisted degradation is more active in BYρ^+^ than BYρ^0^ during chronic proteotoxic stress; this degradation depends on *tom70* and *tim23* on the mitochondrial membrane, *cdc48* in the cytosol, and finally on the proteasome.

AZC treatment led to protein aggregation, as monitored by Hsp104 puncta in BYρ^+^ and BYρ^0^ (Figure 4D). The number of puncta per cell is lower in BYρ^0^ compared to BYρ^+^ (Figure 4D and E). Remarkably, the deletion of *tom70*, which partially prevents the association of misfolded proteins to mitochondria, shows a phenotype like BYρ^0^ (Figure 5L, 5M), suggesting that preventing the mitochondrial association of misfolded proteins allows efficient clearance of aggregates or misfolded proteins. However, we found that degradation of TS22 was not enhanced when prevented from associating with the mitochondrial membrane (in BYρ^0^ or Δ*tom70* or DAmP*tim23*) (Figure 5F and 5I). Therefore, preventing association is necessary but not sufficient to ensure enhanced turnover in BYρ^0^ cells. Taken together we find that the flux through proteasome during proteotoxic stress increased in BYρ^0^, modulated likely by increased expression of ubiquitination pathway enzymes. Concomitantly, degradation associated with mitochondria decreased in this strain compared to BYρ^+^. Differences in the proteome of Δ*tom70* with respect to BYρ^+^ during AZC stress, are in the same direction as shown by BYρ^0^ (Figure 5N). Thus, preventing mitochondrial import or the association of misfolded proteins has the capacity to alter the ability of cells to respond and tolerate misfolding stress.

### Reducing the mitochondrial import components alleviates mitochondrial-associated toxicity

Is the decrease in mitochondria-assisted degradation in BYρ^0^ important for its ability to handle chronic proteotoxic stress? It was clear that chronic expression of TS22 did not cause any fitness defect in BYρ^0^ or BYρ^+^ (Figure 6A, S7A). Consequently, there was no fitness advantage in BYρ^0^ over BYρ^+^ (Figure 6A, S7A). Thus, although TS22 is degraded through mitochondrial association in vivo, it does not lead to a discernible fitness defect. This starkly contrasted with chronic AZC treatment, where BYρ^0^ shows a strong fitness advantage. We hypothesized that the increase in mitochondria-associated misfolded proteins during chronic global proteotoxic stress may be toxic to the cell when a substantial fraction of cellular proteins misfold. Decreasing the association of mitochondria with these misfolded proteins in BYρ^0^ may confer resistance to this stress. Given our hypothesis, we reasoned that blocking the import of misfolded protein in mitochondria would reduce mitochondrial toxicity and hence should increase the fitness of BYρ^+^ itself. To achieve this, we first downregulated *tim23* from parent strain BYρ^+^, effectively inhibiting the import of misfolded proteins into the mitochondria, and examined the effect of this downregulation on the fitness of BYρ^+^ in AZC. Interestingly, the downregulation of *tim23* resulted in a significant increase in the fitness of BY in AZC (Figure 6B). To further substantiate our finding, we deleted or downregulated (as specified) other components of mitochondrial import machinery, encompassing both the outer and inner mitochondrial membrane, and subsequently evaluated the fitness of these deletion strains in the presence of AZC (Figure 6B). As anticipated, a few of the deletions, particularly *tom70,* indeed increased the fitness of BYρ^+^ in AZC. (Figure 6B). AZC tolerance assay with BYρ^+^, BYρ^0^, Δ*tom70*ρ^+^, and Δ*tom70*ρ^0^ (Figure 6C) also validated that *tom70* deletion alone could confer AZC tolerance nearly to the same extent as BYρ^0^. Removal of mtDNA in this background only had a marginal effect on AZC tolerance. We confirmed that these strains harbor mtDNA, and the fitness gain is not due to the loss of mtDNA in these deletion strains (Figure S7B). Additionally, unlike BYρ^+^ (Figure S6A), mitochondrial fragmentation did not increase in BYρ^0^ or Δ*tom70* upon AZC treatment (Figure 6D). Thus mitochondria-associated toxicity decreased when mtDNA or the import receptor was deleted.

**Figure 6.**
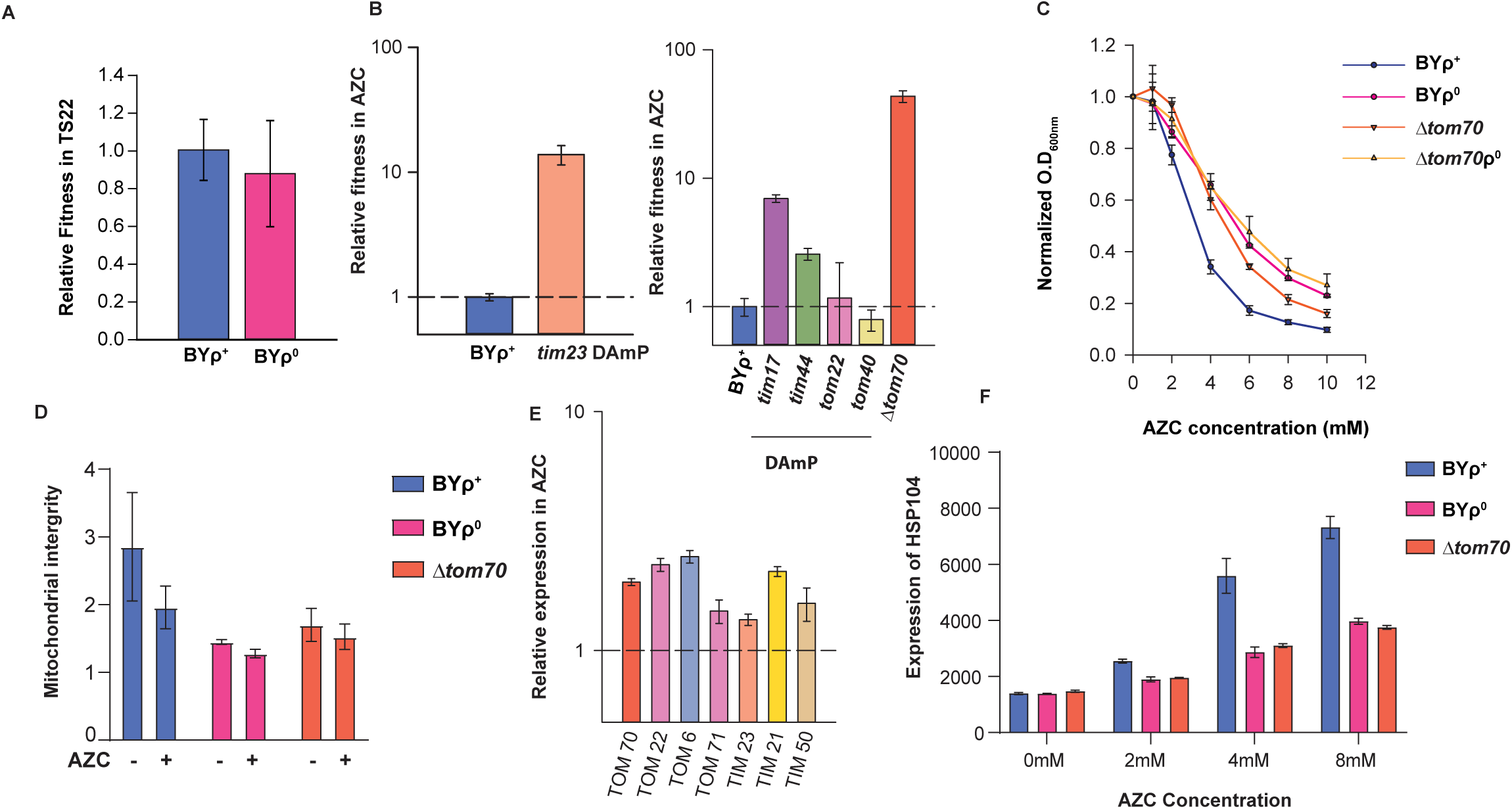
Reducing the mitochondrial import components alleviates toxicity. A) Relative fitness of BYρ^0^ and BYρ^+^ in presence of single misfolded protein TS22. BYρ^0^ and BYρ^+^ cells expressing TS22 were mixed with TDH2-GFP cells expressing TS22 and outcompeting population was monitored by flow cytometry. Fitness of TS22 is normalized wrt cells expressing WT Nat-R. B) Relative fitness of cells with downregulation of *tim23* (left panel) and other import machinery components (right panel) in AZC at 48 hours. C) Increased tolerance of Δ*tom70* as compared to BYρ^+^ in AZC. D) Mitochondrial fragmentation as measured by shape analysis in different strains containing mitochondrial mCherry plasmid, with and without 4mM AZC treatment. E) Relative expression of mitochondrial import components upon treatment with 4mM AZC. GFP tagged strains of import components are utilized for monitoring the changes in expression by flow cytometry. F) Relative expression of HSP104 in BYρ^0^, BYρ^+^ and Δ*tom70* upon treatment with increasing concentration of AZC at 30°C.

Notably, we found the mitochondrial import machinery to be upregulated in BYρ^+^ in AZC (Figure 6E), suggesting that misfolding signals mitochondrial import activation which may exacerbate this unanticipated toxicity. To confirm that the gain in fitness in the import channel deletion strains during AZC treatment, is linked to a decrease in proteotoxicity in them, we monitored the canonical proteotoxic stress response pathway in BYρ^0^ and Δ*tom70* cells. These pathways were less activated in both BYρ^0^ and Δ*tom70* (Figure 6F, S7C) than in BYρ^+^ in AZC. Thus decrease in mistargeting of misfolded proteins to the mitochondrial surface decreases proteotoxic stress to cells and increases tolerance towards proteotoxicity.

## DISCUSSION

We identified two key factors that enable BYρ^0^ cells to tolerate proteotoxicity. The first factor is their ability to prevent misfolded proteins from associating with the mitochondrial surface, thereby reducing toxicity. The second factor is their capacity for the rapid degradation of misfolded proteins. While these two abilities are interconnected, the enhanced degradation observed in BYρ^0^ is likely a cellular response to misfolding stress. Our findings suggest that mitochondria are active in maintaining cytosolic proteostasis, particularly in clearing misfolded proteins when the misfolding stress is not global. However, this protective mechanism may become detrimental during a global misfolding event, as the increased load of misfolded proteins on the mitochondrial surface could lead to toxicity. Understanding this process and the critical tipping point where it shifts from being protective to being harmful, may have implications for many instances where proteostasis control is envisaged.

### Mitochondria-associated degradation of cytosolic misfolded proteins

In this study, we found that misfolded proteins are mistargeted to the surface of mitochondria and undergo proteasome-mediated degradation. While this is reminiscent of MAD^37,53–55^ (or mitoTAD^36^) pathways, there is a subtle difference. While these reported pathways take care of misfolding of mitochondrially targeted proteins (sometimes ER proteins) that are mistargeted or clogged in the import channels on mitochondria, the pathway we discuss, is involved in clearing cytosolic misfolded proteins. This, while being like MAGIC^49^ in its essence, is different since the degradation in MAGIC is proposed to occur inside mitochondrial compartments. Our proposed pathway clears protein using proteasome and the canonical clearance pathway that takes care of clogged mitochondrial import channels^36^. Unlike MAGIC, this pathway is operational in clearing misfolded proteins without a heat shock. Interestingly, this degradation pathway efficiently clears a single misfolded protein species without taxing cellular fitness. During global misfolding events, a larger dose of misfolded protein on the surface of mitochondria decreases cellular fitness. There was a hint of this phenomenon in a recent report^50^. However, the authors focused on protein aggregation mediated by Tom70 and its substrates. Using multiple genetic models, we show that the toxicity due to misfolding is Tom70-dependent and dependent on other factors regulating mitochondrial import. Although we could not demonstrate how this decreased cellular fitness, we ruled out one crucial factor: a decrease in the import capacity of mitochondria due to the channels being titrated out with misfolded proteins. Decreasing import capacity during misfolding stress decreases the association of misfolded proteins on the mitochondrial surface, alleviating proteotoxicity.

The association of misfolded proteins on mitochondria has been reported earlier for huntingtin with CAG repeat expansion ^56,57^, Aβ and model misfolded protein like DM-Luc^49,52^. This prompted the research community to search for evidence of import-channel block as the primary mechanism of toxicity in dominant misfolding diseases^58^. Although Huntingtin interacted with Tim23, it was reported not to enter the mitochondrial compartment in vivo^59^. If our work holds in higher eukaryotes, we speculate that the toxicity stems from the association of the misfolded protein with mitochondria and not their import. Similarly, from our studies, we hypothesize that the association is toxic in addition to their potential role in blocking import^57^.

While proteosomes can receive substrates routed through mitochondria, without this, ρ^0^ cells can clear out misfolded proteins more efficiently during misfolding stress. This indicates two parts to the problem of solving fitness defects during proteotoxic stress. First, decrease in the association of misfolded proteins with mitochondria. Second, better clearance of the misfolded proteins during proteotoxic stress even when the quantity of proteasome is similar in ρ^0^ and ρ^+^ cells. We find ubiquitinating enzymes upregulated, indicating that processes upstream of degradation may be upregulated in these strains. While this is not resolved completely in this manuscript, we hypothesize the upregulation of the ubiquitination machinery in ρ^0^ cells during misfolding stress to be essential for this (Figure 4J). Additionally, low Hsp90 in the ρ^0^ cells during misfolding stress could be helpful in protein degradation. Hsp90 is known to act like a sink for misfolded proteins that take over unfolded proteins from Hsp70^60^. Hsp90 also acts like a reported brake on Hsp70 activity by aiding proteins to fold^60^. Given this, we speculate that low Hsp90 allows the substrate to stably associate with Hsp70s of the cytosol, allowing them to be channeled for degradation through proteasome more efficiently than in ρ^+^ cells where Hsp90 is upregulated during misfolding stress.

### Limitations of this study

The work is primarily done in a single-cell eukaryotic model, so more work is needed to confirm the conservation of this mechanism. Additionally, we have used different grades of misfolding mutants of the same protein (Nat-R). This is unlikely to be a particular case, but further studies using multiple proteins, especially endogenous ones, will be required to understand the generality of the process. Although we see the global association of ubiquitinated proteins with mitochondria during misfolding stress, we do not know if all misfolded proteins follow the same degradation path. Therefore, the rules governing the association of misfolded proteins with mitochondria must be delineated.

### Summary and Implications

While the stage is too nascent to comment on the implication of the process, it is essential to note that the association of misfolded proteins with mitochondria has been repeatedly reported in the literature, but with various downstream pathways being proposed. Here we start with an unbiased approach to reach a surprising conclusion; mitochondria uses quality control systems to take care of protein import and cater to cytosolic protein quality control. A small dose of misfolded proteins can be efficiently handled by the organelle, but during a proteome-wise misfolding event, the same may be a fitness cost for the cell. We have also noticed that this may have further ramifications in protein folding if the association of misfolded proteins is mediated by import channels; repeated interaction in the presence of mitochondrial potential also has the capacity, in theory, to disentangle protein aggregates or remove kinetic traps by unfolding misfolded proteins. We have seen that TS22, the temperature-sensitive protein, is more active in BYρ^+^ cells with active mitochondrial association than in BYρ^0^ cells lacking this association (Figure 4A). This holds true for tim23 DAmP which shows lower degradation of TS22, yet lower activity like in BYρ^0^ (Figure S7E, Figure 4A). Thus, suggesting that the mitochondrial association, while having a role in degradation, may have implications in folding as well. Whether this process occurs and complements the mitochondria-assisted degradation of misfolded proteins would be essential to dissect in the context of cytosolic proteostasis.

## METHODS

### Yeast strains, plasmids, and growth conditions

*Saccharomyces*L*cerevisiae* strains used in this study (Table S1). Yeast cells were cultivated on rich YPD medium containing 1% (w/v) yeast extract, 2% (w/v) peptone and 2% (w/v) dextrose or non-fermentative carbon sources (2% glycerol, 2% ethanol). When using deletion or strains transformed with plasmids, synthetic drop-out media was used for selection. Strains were grown at 30[ at 200 rpm unless mentioned otherwise. Cells were inoculated in YPD at 0.2 O.D_600_ from an overnight primary culture and grown to mid-log phase before harvesting for further experiments.

### Generation of thermotolerant *Saccharomyces cerevisiae* strains through Adaptive Laboratory Evolution

*Saccharomyces cerevisiae* strain BY4741 (MATa his3Δ1 leu2Δ0 lys2Δ0 ura3Δ0) was used as the parent strain for evolution. Single colonies were inoculated in 5 ml of rich medium (YPD) and were grown overnight at 30°C (permissive temperature). Saturated primary cultures were then re-inoculated in deep multi-well plates at an O.D_600_ of 0.2 and shifted to a growth temperature of 40°C at 100 rpm in a water bath. A total of 29 such parallel evolutions were conducted for 600 generations at 40°C. Passages were performed every 16-20 hours where 1% of the inoculum from each well was transferred to a new deep well plate containing around 400[l of fresh media. Glycerol stocks were maintained at -80°C after every 100 generations, and the growth phenotypes and contaminations of evolved strains were monitored. Same protocol was followed for adaptive laboratory evolution of BY4741, W303 and RM11-1a for 100 generations, where glycerol stocks were made after every 10 generations.

### Quantification And Statistical Analysis

Student’s t-test and R package for non-linear regression was used for statistical analysis. Mann-Whitney-U test was used for pathway specific analysis using Wilcox package in R.

## DATA AND SOFTWARE AVAILABILITY

All data are provided in the manuscript. Genome sequencing files are under bioproject accession PRJNA1126056

Please see the supplemental information section for detailed materials and methods.

## ACKNOWLEDGEMENTS

We acknowledge Patrick De’Silva (IISc Bangalore) for sharing antibody against yeast Tim23, Koyeli Mapa (SNU) for her help with mitochondria related work. We acknowledge the imaging facility of CSIR-IGIB for microscopy, and the IT of CSIR-IGIB for computational facility. We acknowledge Mohammad Aaquib for DM-MBP purification and labelling. K.C. acknowledges funding from the CSIR Empower grant (MLP2103) which initiated the work, and funding from DBT-Wellcome Trust India-Alliance grant IA/S/21/1/505587 which subsequently funded the work. Z.Z and S.B acknowledges CSIR-IGIB; D.P.D acknowledges DBT; A.S and S.R acknowledge UGC, K.P and S.K acknowledge CSIR for fellowship support.

## AUTHOR CONTRIBUTION

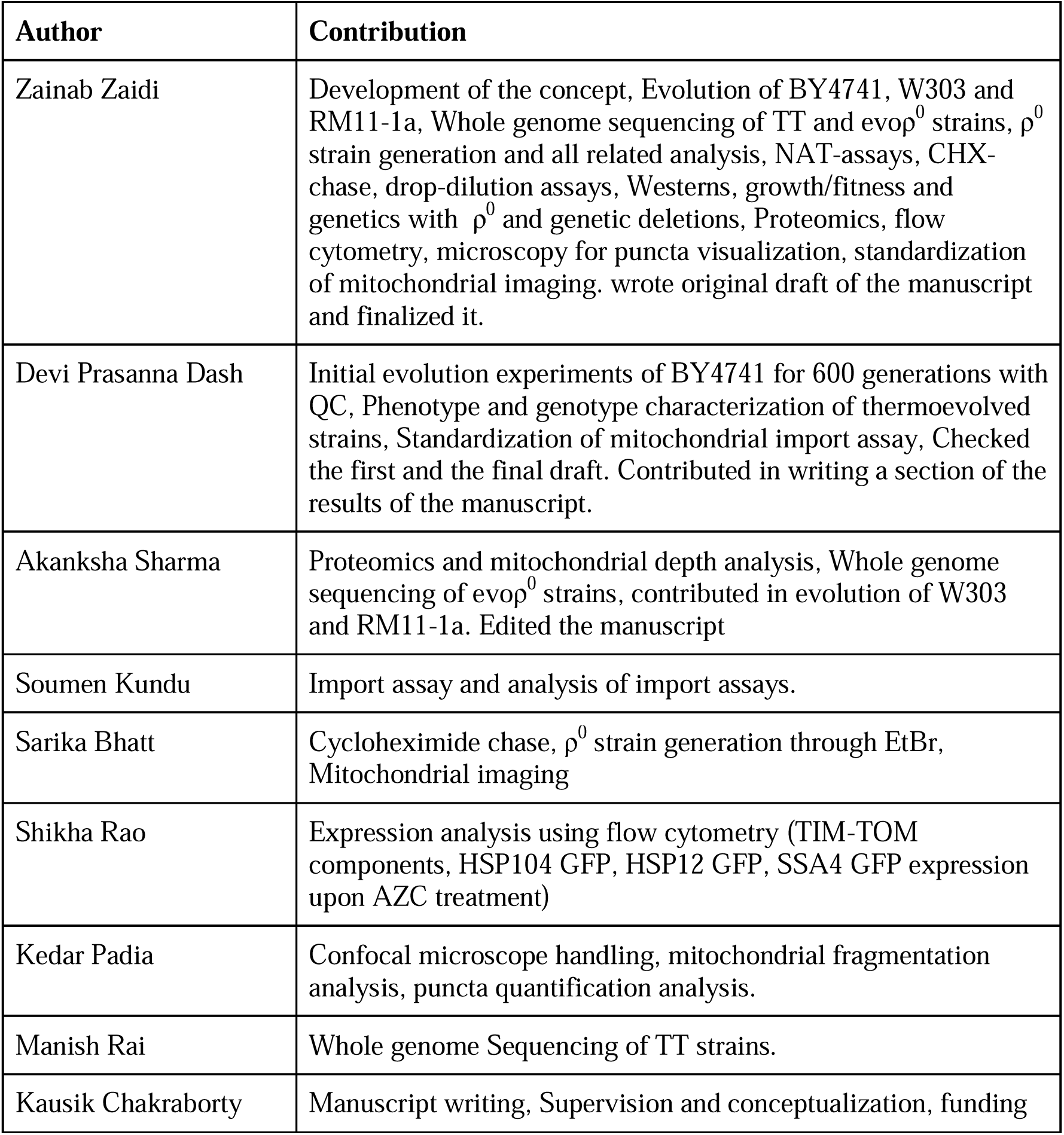
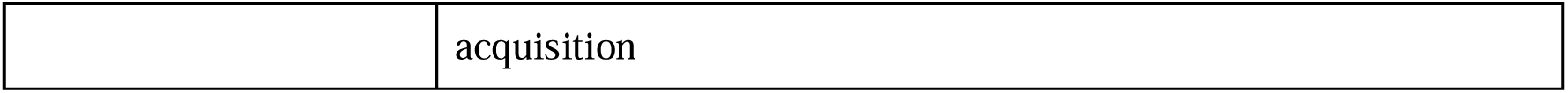

## REFERENCES

1. Hartl, F. U. Molecular chaperones in cellular protein folding. Nature 381, 571–580 (1996).

2. Balch, W. E., Morimoto, R. I., Dillin, A. & Kelly, J. W. Adapting proteostasis for disease intervention. Science 319, (2008).

3. Hipp, M. S., Kasturi, P. & Hartl, F. U. The proteostasis network and its decline in ageing. Nat. Rev. Mol. Cell Biol. 20, 421–435 (2019).

4. Chen, B., Retzlaff, M., Roos, T. & Frydman, J. Cellular Strategies of Protein Quality Control. Cold Spring Harb. Perspect. Biol. 3, (2011).

5. Ciechanover, A. The ubiquitin-proteasome pathway: on protein death and cell life. EMBO J. 17, 7151 (1998).

6. Dobson, C. M. Protein folding and misfolding. Nature 426, (2003).

7. Valastyan, J. S. & Lindquist, S. Mechanisms of protein-folding diseases at a glance. Dis. Model. Mech. 7, (2014).

8. Chiti, F. & Dobson, C. M. Protein Misfolding, Amyloid Formation, and Human Disease: A Summary of Progress Over the Last Decade. Annu. Rev. Biochem. 86, (2017).

9. Verma, K., Verma, M., Chaphalkar, A. & Chakraborty, K. Recent advances in understanding the role of proteostasis. Faculty Reviews 10, (2021).

10. Krobitsch, S. & Lindquist, S. Aggregation of huntingtin in yeast varies with the length of the polyglutamine expansion and the expression of chaperone proteins. Proceedings of the National Academy of Sciences 97, 1589–1594 (2000).

11. Walsh, D. M. et al. Naturally secreted oligomers of amyloid β protein potently inhibit hippocampal long-term potentiation in vivo. Nature 416, 535–539 (2002).

12. Winner, B. et al. In vivo demonstration that α-synuclein oligomers are toxic. Proceedings of the National Academy of Sciences 108, 4194–4199 (2011).

13. Behrends, C. et al. Chaperonin TRiC Promotes the Assembly of polyQ Expansion Proteins into Nontoxic Oligomers. Mol. Cell 23, 887–897 (2006).

14. Yu, A. et al. Protein aggregation can inhibit clathrin-mediated endocytosis by chaperone competition. Proceedings of the National Academy of Sciences 111, E1481–E1490 (2014).

15. Park, S.-H. et al. PolyQ Proteins Interfere with Nuclear Degradation of Cytosolic Proteins by Sequestering the Sis1p Chaperone. Cell 154, 134–145 (2013).

16. de Groot, N. S. et al. The fitness cost and benefit of phaselseparated protein deposits. Mol. Syst. Biol. 15, (2019).

17. Olzscha, H. et al. Amyloid-like aggregates sequester numerous metastable proteins with essential cellular functions. Cell 144, (2011).

18. Lashuel, H. A., Hartley, D., Petre, B. M., Walz, T. & Lansbury, P. T. Amyloid pores from pathogenic mutations. Nature 418, 291–291 (2002).

19. Yablonska, S. et al. Mutant huntingtin disrupts mitochondrial proteostasis by interacting with TIM23. Proceedings of the National Academy of Sciences 116, 16593–16602 (2019).

20. Chronic exposure to subllethal betalamyloid (Aβ) inhibits the import of nuclearlencoded proteins to mitochondria in differentiated PC12 cells*. 10.1111/j.1471-4159.2007.04907.x.

21. Duennwald, M. L. & Lindquist, S. Impaired ERAD and ER stress are early and specific events in polyglutamine toxicity. Genes Dev. 22, 3308–3319 (2008).

22. Park, S. K. et al. Overexpression of the essential Sis1 chaperone reduces TDP-43 effects on toxicity and proteolysis. PLoS Genet. 13, (2017).

23. Sakahira, H., Breuer, P., Hayer-Hartl, M. K. & Hartl, F. U. Molecular chaperones as modulators of polyglutamine protein aggregation and toxicity. Proceedings of the National Academy of Sciences 99, 16412–16418 (2002).

24. Catarina Silva, M., et al. A Genetic Screening Strategy Identifies Novel Regulators of the Proteostasis Network. PLoS Genet. 7, e1002438 (2011).

25. Willingham, S., Outeiro, T. F., DeVit, M. J., Lindquist, S. L. & Muchowski, P. J. Yeast Genes That Enhance the Toxicity of a Mutant Huntingtin Fragment or α-Synuclein. Science (2003) doi:10.1126/science.1090389.

26. Khurana, V. & Lindquist, S. Modelling neurodegeneration in Saccharomyces cerevisiae: why cook with baker’s yeast? Nat. Rev. Neurosci. 11, 436–449 (2010).

27. Kaiser, C. J. O. et al. A network of genes connects polyglutamine toxicity to ploidy control in yeast. Nat. Commun. 4, 1–11 (2013).

28. Nollen, E. A. A. et al. Genome-wide RNA interference screen identifies previously undescribed regulators of polyglutamine aggregation. Proceedings of the National Academy of Sciences 101, 6403– 6408 (2004).

29. Giorgini, F., Guidetti, P., Nguyen, Q., Bennett, S. C. & Muchowski, P. J. A genomic screen in yeast implicates kynurenine 3-monooxygenase as a therapeutic target for Huntington’s disease. Nat. Genet. 37, 526 (2005).

30. Caspeta, L. et al. Altered sterol composition renders yeast thermotolerant. Science (2014) doi:10.1126/science.1258137.

31. Caspeta, L., Chen, Y. & Nielsen, J. Thermotolerant yeasts selected by adaptive evolution express heat stress response at 30 °C. Sci. Rep. 6, 1–9 (2016).

32. Yona, A. H. et al. Chromosomal duplication is a transient evolutionary solution to stress. Proc. Natl. Acad. Sci. U. S. A. 109, (2012).

33. Rudolph, B., Gebendorfer, K. M., Buchner, J. & Winter, J. Evolution of Escherichia coli for Growth at High Temperatures. J. Biol. Chem. 285, 19029 (2010).

34. Tenaillon, O. et al. The Molecular Diversity of Adaptive Convergence. Science (2012) doi:10.1126/science.1212986.

35. Ghosh, K. & Dill, K. Cellular proteomes have broad distributions of protein stability. Biophys. J. 99, (2010).

36. Mårtensson, C. U. et al. Mitochondrial protein translocation-associated degradation. Nature 569, 679– 683 (2019).

37. Regulating mitochondrial outer membrane proteins by ubiquitination and proteasomal degradation. Curr. Opin. Cell Biol. 23, 476–482 (2011).

38. Laboratory adaptive evolution of thermotolerance is linked to the evolution of a robust proteostasis in S. cerevisiae. Paperpile https://paperpile.com/app/p/9f1fae88-9a34-0aa2-8916-902f4b124e5f.

39. Huh, W. K. et al. Global analysis of protein localization in budding yeast. Nature 425, (2003).

40. Laboratory adaptive evolution of thermotolerance is linked to the evolution of a robust proteostasis in S. cerevisiae. Paperpile https://paperpile.com/app/p/9f1fae88-9a34-0aa2-8916-902f4b124e5f.

41. Jarolim, S. et al. Saccharomyces cerevisiae Genes Involved in Survival of Heat Shock. G3: Genes|Genomes|Genetics 3, 2321 (2013).

42. Huang, C. J., Lu, M. Y., Chang, Y. W. & Li, W. H. Experimental Evolution of Yeast for High-Temperature Tolerance. Mol. Biol. Evol. 35, (2018).

43. Baruffini, E., Lodi, T., Dallabona, C. & Foury, F. A Single Nucleotide Polymorphism in the DNA Polymerase Gamma Gene of Saccharomyces cerevisiae Laboratory Strains Is Responsible for Increased Mitochondrial DNA Mutability. Genetics 177, 1227 (2007).

44. Replacement of proline by azetidine-2-carboxylic acid during biosynthesis of protein. Biochim. Biophys. Acta 71, 459–461 (1963).

45. Trotter, E. W. et al. Misfolded Proteins Are Competent to Mediate a Subset of the Responses to Heat Shock in Saccharomyces cerevisiae *. J. Biol. Chem. 277, 44817–44825 (2002).

46. Retrograde regulation of multidrug resistance in Saccharomyces cerevisiae. Gene 354, 15–21 (2005).

47. Ghosh, A. et al. Cellular responses to proteostasis perturbations reveal non-optimal feedback in chaperone networks. Cell. Mol. Life Sci. 76, (2019).

48. Kaganovich, D., Kopito, R. & Frydman, J. Misfolded proteins partition between two distinct quality control compartments. Nature 454, 1088 (2008).

49. Ruan, L. et al. Cytosolic proteostasis through importing of misfolded proteins into mitochondria. Nature 543, 443–446 (2017).

50. Wang, Y. et al. Metabolic regulation of misfolded protein import into mitochondria. (2024) doi:10.7554/eLife.87518.

51. Wang, J. D., Michelitsch, M. D. & Weissman, J. S. GroEL-GroES-mediated protein folding requires an intact central cavity. Proceedings of the National Academy of Sciences 95, 12163–12168 (1998).

52. Liu, Q. et al. Nascent mitochondrial proteins initiate the localized condensation of cytosolic protein aggregates on the mitochondrial surface. Proceedings of the National Academy of Sciences 120, e2300475120 (2023).

53. Xu, S., Peng, G., Wang, Y., Fang, S. & Karbowski, M. The AAA-ATPase p97 is essential for outer mitochondrial membrane protein turnover. Mol. Biol. Cell (2010) doi:10.1091/mbc.e10-09-0748.

54. Website. 10.1083/jcb.201510098.

55. Website. https://rupress.org/jcb/article/213/1/49/38547/Doa1-targets-ubiquitinated-substrates-for.

56. Orr, A. L. et al. N-Terminal Mutant Huntingtin Associates with Mitochondria and Impairs Mitochondrial Trafficking. J. Neurosci. 28, 2783–2792 (2008).

57. Yano, H. et al. Inhibition of mitochondrial protein import by mutant huntingtin. Nat. Neurosci. 17, 822– 831 (2014).

58. Needs, H. I., Wilkinson, K. A., Henley, J. M. & Collinson, I. Aggregation-prone Tau impairs mitochondrial import, which affects organelle morphology and neuronal complexity. J. Cell Sci. 136, (2023).

59. Hamilton, J., Brustovetsky, T., Khanna, R. & Brustovetsky, N. Mutant huntingtin does not cross the mitochondrial outer membrane. Hum. Mol. Genet. 29, (2020).

60. Morán, L. T., Kityk, R., Mayer, M. P. & Rüdiger, S. G. D. Hsp90 Breaks the Deadlock of the Hsp70 Chaperone System. Mol. Cell 70, (2018).

