## Supplemental methods and figure legends for "Fitness defects due to cytosolic protein misfolding in *S. cerevisiae* can be alleviated by decreasing mitochondrial protein import capacity"

### **Yeast strains, plasmids, and growth conditions**

### *Saccharomyces cerevisiae* strains used in this study are given in key resource table. Yeast cells were cultivated on rich YPD medium containing 1% (w/v) yeast extract, 2% (w/v) peptone and 2% (w/v) dextrose or non-fermentative carbon sources (2% glycerol, 2% ethanol). When using deletion or strains transformed with plasmids, synthetic drop-out media was used for selection. Strains were grown at 30℃ at 200 rpm unless mentioned otherwise. Cells were inoculated in YPD at 0.2 O.D_600_ from an overnight primary culture and grown to mid-log phase before harvesting for further experiments.

### **Generation of thermotolerant *Saccharomyces cerevisiae* strains through Adaptive Laboratory Evolution**

*Saccharomyces cerevisiae* strain BY4741 (MATa his3Δ1 leu2Δ0 lys2Δ0 ura3Δ0) was used as the parent strain for evolution. Single colonies were inoculated in 5 ml of rich medium (YPD) and were grown overnight at 30°C (permissive temperature). Saturated primary cultures were then re-inoculated in deep multi-well plates at an O.D_600_ of 0.2 and shifted to a growth temperature of 40°C at 100 rpm in a water bath. A total of 29 such parallel evolutions were conducted for 600 generations at 40°C. Passages were performed every 16-20 hours where 1% of the inoculum from each well was transferred to a new deep well plate containing around 400𝜇l of fresh media. Glycerol stocks were maintained at -80°C after every 100 generations, and the growth phenotypes and contaminations of evolved strains were monitored. Same protocol was followed for adaptive laboratory evolution of BY4741, W303 and RM11-1a for 100 generations, where glycerol stocks were made after every 10 generations.

**Growth assay**

Primary cultures were grown overnight (~18 hours) at 30° C. The saturated cultures were re-inoculated in fresh YPD medium at an O.D_600_ of 0.1 in honeycomb plate. Growth assay for all TT strains were done at 30⁰C and 40⁰C for 24 hours.

**Drop Dilution Assay**

Single colonies were inoculated in 3 ml of YPD medium and were grown overnight at 30°C as primary cultures. Secondary cultures were inoculated at O.D_600_ of 0.2 and were grown till log-phase. Cells were pelleted, washed and resuspended in Autoclaved MQ to a final concentration of O.D_600_ =0.4/ml. Serial 1:10 dilutions were prepared and 4 ul of each dilution were spotted on YP agar plates containing either 2% of fermentable (glucose), 2% of non-fermentable carbon source (glycerol, lactate, ethanol) or YPD agar plates with L-Azetidine-2-carboxylic acid (AZC). Plates were incubated at 30°C or 40°C (as specified) for 2 days and photographed.

**Competitive Fitness Assay**

Single colonies of each strain were inoculated in 400𝜇l of YPD medium in a deep well plate along with TDH2-GFP from Yeast-GFP Clone collection. The cultures were grown overnight at 30°C on 200rpm as primary cultures. The assay was set up by mixing an equal O.D_600_ of cells of the control TDH2-GFP and thermotolerant evolved strain in a well. The cultures were grown and passaged after every 12 hours and the proportion of each population was measured by Flow cytometer BD LSR II after every 24 hours or as indicated otherwise. To evaluate thermotolerance in the evolved strains of S. cerevisiae, the cultures were incubated in YPD medium at both 30°C and 40°C. For fitness assessment in the presence of AZC, a concentration of 4 mM AZC was used, and the cultures were grown at 30°C in an incubator shaking at 200 rpm.

**Measurement of Oxygen Consumption Rate**

Yeast strains were grown in YPD medium, and secondary culture were inoculated at an OD600 of 0.2 and grown till log-phase at 30°C. Cells were then diluted with media to an O.D_600_ equivalent to 1.0 and used to measure the oxygen consumption rate using Oxygraph (Oroboros) instrument. Initially the chambers were filled with fresh YPD medium to establish the standard oxygen levels in the chamber, followed by the addition of cells to measure their oxygen consumption. The parent strain BY4741 was used as a control to assess difference in oxygen consumption rate.

**quantitative Polymerase Chain Reaction (qPCR)**

Genomic DNA was extracted using the LiAc-SDS method, and 50 ng of DNA was used as input for each reaction. The real-time quantitative PCR (qPCR) assays were performed using the thermocycler. Each 10 μl reaction mixture contained 1 μM of forward and reverse primers, 50 ng of genomic DNA, and 1X SYBR mix (Takara) with the final volume adjusted using NFW. PCR conditions were 95 °C for 5 min followed by 40 cycles of 95 °C for 30 s and 55 °C for 30 s and 72 °C for 1 min and a final 10 min extension at 72 °C. Primers targeting the mtCOX3 gene were used for the mitochondrial gene, while the nHsp31 gene served as a control. The abundance of mtCOX3 was quantified by normalizing the Ct values of the mtCOX3 gene in each strain to their respective nHsp31 control, followed by further normalization wrt BY4741.

**Cycloheximide Chase Assay**

From overnight-grown primary cultures, secondary cultures were inoculated at 0.2 O.D_600_ in 50ml of synthetic-defined uracil drop-out media (SD-ura). The cultures were allowed to grow till they reach O.D_600nm_ of 0.4-0.6. Once reached, the cultures were harvested and resuspended in 10ml of spent media, making a high-density culture. In the high-density culture, cycloheximide was added to a final concentration of 40𝜇g/ml and the translation was arrested. Post-translational arrest, 2ml aliquotes were collected at 0, 30, 60, and 90 minutes (or as specified). The cells were harvested, washed and snap frozen. The protein was isolated by the alkaline lysis method, followed by protein estimation by Pierce^TM^ BCA kit (Thermo Scientific, Cat no-23225) and western blotting.

**Western Blotting**

Secondary culture was inoculated at ~ 0.2 O.D_600_ and grown at 30 °C; 200 rpm. The cultures were pelleted and protein was isolated by alkaline lysis method, followed by protein estimation by Pierce^TM^ BCA kit (Thermo Scientific, Cat no-23225). For SDS-PAGE and western blotting, 30 μg of protein was loaded and transferred onto a nitrocellulose membrane (Millipore; HATF00010). Following transfer, the membrane was stained with ponceau stain to procure the image (loading control). The membrane was blocked with 2% BSA in TBST for 90 minutes and incubated with the desired primary antibody overnight at 4°C. The next day, the membrane was washed three times with TBST (15 minutes each) and incubated with an HRP-conjugated secondary antibody (anti-rabbit or anti-mouse) for 2 hours at room temperature. Following secondary antibody incubation, the membrane was washed again to remove non-specific binding. Blots were developed using Immobilon Western Chemiluminescent HRP substrate (Millipore, WBKLS0500) and imaged using the SYNGENE gel documentation system.

**Growth Assay in presence of antibiotic**

From an overnight primary culture, the cells were grown in the presence of varying concentrations of nourseothricin (Jena Biosciences) ranging from 0𝜇g/ml to 100𝜇g/ml at 30°C, 200 rpm. 400 ul of culture (initial O.D_600_ being 0.05-0.1) was kept per well and grown at 30 °C; 200 rpm for 16 h. Growth readings were taken on a multi-plate reader (TECAN infinite 200 pro).

Growth assay was performed similarly with AZC (L-Azetidine-2-carboxylic acid; A0760 Sigma) with concentration ranging from 0-10mM.

**Generation of ρ^0^ strain by Ethidium Bromide treatment**

For the induction of rho mutants, cells were grown at 30 °C in SC medium until reaching an O.D_600_ of 0.5-0.6, followed by treatment with 25 μg/ml of ethidium bromide ([Slonimski et al. 1968](https://www.sciencedirect.com/science/article/pii/S0944501313000979#bib0210)) for 48 hours. After treatment, the cells were diluted in sterile water and plated on YPD agar for isolation of petite colonies. The rho status of the colonies was confirmed by their inability to grow on a non-fermentative carbon source (YP+2% glycerol agar) and by PCR analysis check for the absence of the Cox-3 gene sequence.

**Confocal microscopy**

Yeast cells were grown at 30°C in YPD or YPD medium containing 4mM AZC until they reached an O.D_600_ of 0.5-0.6. Cells were harvested, washed once with 1X PBS and resuspended in 1X PBS. To prepare the slides for confocal microscopy, 200-400𝜇l of fresh concanavalin A solution (0.5mg/ml) was coated on each slide (single slide or chambered slide) and incubated for 30 minutes followed by washing with 1X PBS thrice to remove excess concanavalin A. To the Con A coated slides, around 500𝜇l of resuspended cells were added and incubated for 30 minutes. The slides were washed with 1X PBS thrice to remove unbound excess cells and to achieve optimum density. The cells were covered with a cover slip (thickness aspect #1.5) and a mounting agent and imaged in a confocal microscope. In the case of chambered slides, around 1-2ml of PBS or media was added to the chambers.

**Imaging of yeast mitochondria**

Cells were transformed with plasmid containing mitochondrial targeted mCherry and selected on SD-Ura plates. For microscopy, cells were grown in synthetic defined (SD-Ura medium) till the OD600 of 0.5 and coated on concanavalin A slides as discussed previously. Imaging was done in Leica TCS SP8 confocal microscope.

**Whole Genome Sequencing**

The genomic DNA of the strains was isolated using the glass bead method. Before proceeding with library preparation, the quality and concentration of the genomic DNA were assessed by agarose gel and Qubit high-sensitivity DNA kit respectively. The library preparation was performed using the Illumina Nextera XT library preparation kit (15031942), and indexing was carried out using the dual indexing kit (Nextera XT Index kit v2). The pooled gDNA library was sequenced on the Illumina Hiseq 2500 platform with 300*2 bp paired-end reads and a target coverage of approximately 40X per sample. FastQ files were generated using the Illumina bcl2 fastq Conversion software. The quality of the reads was assessed using FastQC, and low-quality reads were filtered out using trimmomatic (Bolger et al., 2014). Read alignment to the reference genome of yeast was performed using BWA (Li and Durbin, 2010). SAM tools was utilized for the post-processing of the BAM file and the removal of duplicate reads (Li et al., 2009). Mitochondrial DNA depth was determined using SAM tools and represented using R packages.

**Proteomics**

Protein was isolated from the desired cultures using the alkaline lysis method. Protein concentration was determined using Pierce^TM^ BCA kit (Thermo Scientific, Cat no-23225) and 100𝜇g of protein was aliquoted in a fresh microcentrifuge tube. Only unautoclaved plasticware was used from this step onward. To the aliquoted protein lysate, around 600𝜇l of chilled acetone was added, mixed thoroughly and kept overnight at -20°C for precipitation of protein. The next day, the samples were spun down at high speed for 10-15 minutes and the pellet was air dried for around 5 minutes. To the dried protein pellet, 40𝜇l of Tris Urea buffer of pH 8.5 was added and the protein concentration was determined using Bradford assay (Sigma, B6916). 20𝜇g of protein was aliquoted in a fresh microcentrifuge tube for further processing. The protein was treated with 2𝜇l of DTT solution (25mM) at 56°C for 30 minutes followed by treatment with 1𝜇l of Iodoacetamide solution (55mM) in the dark for 15 minutes. This was diluted by adding 7 volumes (7 times the protein lysate volume) of 50mM Tris buffer, pH 8 to dilute the urea present in the sample. To this, 2𝜇l of trypsin (1𝜇g/𝜇l, used in 1:20 ratio) was added and mixed thoroughly and incubated at 37°C for 16 -18 hours on shaking in a dry bath. The reaction was quenched by adding 1𝜇l 0.1% formic acid and the sample was dried in a SpeedVac vacuum concentrator in V-Aq mode at 30°C. The pellet can be stored at -20°C or directly processed for desalting. In further processing, the pellet was resuspended in 15 𝜇l of 0.1% formic acid and vortexed properly, followed by a brief spin. For desalting, zip tips were used in the following sequence: 10 times in 100% Acetonitrile (ACN), 10 times in 0.1% formic acid, 15 times in sample, 2 times in 5% methanol and lastly 15 times pipette up and down in 50𝜇l of elution buffer having 70% ACN and 0.1% formic acid. After desalting, the eluted peptide solution was vacuum dried using SpeedVac vacuum concentrator in V-Aq mode at 30°C for 2 hours. The lyophilized peptides were resuspended in 15𝜇l of 0.1% formic acid, vortexed and spun down at 10000 rpm for 5 minutes. The fractionated peptides were analyzed on a quadrupole-TOF hybrid mass spectrometer (TripleTOF 6600, AB Sciex, USA) coupled to a nano-LC system (Eksigent NanoLC-425) in SWATH (relative quantitation) mode. For this, 5𝜇g of desalted peptides were loaded onto a trap column (chromXP C18CL 5𝜇m 120 Å, Eksigent) at a flow rate of 5𝜇l/minute using Buffer A (water with 0.1% formic acid) and Buffer B (acetonitrile with 0.1% formic acid). The data was analyzed using Spectraunaut software and represented using R packages.

**Isolation of mitochondria for import assay**

From an overnight-grown primary culture, secondary inoculations were done in 50-100 ml of YPD (or YPD with 4 mM AZC, as specified) and grown at 30°C, 200 rpm till saturated. The saturated culture was harvested by centrifugation at 2500 x g for 10 minutes at 4°C, followed by washing with AMQ. The pellet was then resuspended in 5 ml DTT buffer (100 mM Tris-SO_4_, 10 mM DTT, pH 9.4). The cell suspension was incubated for 30 minutes at 30°C with gentle shaking and was spun down at 2500 x g for 10 minutes at 4°C. The pellet was resuspended in 5 ml of 1.2 M sorbitol buffer (1.2 M sorbitol, 20 mM Potassium phosphate, pH 7.4). To this, zymolase (2.5 mg/gm of pellet weight) was added and incubated for 40 minutes at 30°C to digest the cell wall. To monitor the formation of spheroplasts, the cell suspension was diluted in AMQ and sorbitol buffer separately and their O.D_600nm_ was measured. The zymolase treatment was stopped if the O.D_600nm_ of AMQ dilution was 10-20% of the sorbitol dilution, which denotes the bursting of spheroplast in water (clear solution). The spheroplasts were centrifuged at 2500 x g for 5 minutes at 4°C followed by resuspension in 5 ml of homogenization buffer (0.6 M sorbitol, 20 mM HEPES, 1 mM EDTA, pH 7.4) and homogenization in a Dounce-Homogenizer (50-60 times). After this, the cell remnants and unopened cells were pelleted by centrifugation at 2500 x g for 5 minutes at 4°C twice. The supernatant was transferred in an autoclaved Oakridge tube and centrifuged at high speed 18,000 x g for 45 minutes at 4°C. The cytosolic fraction was collected in a 15 ml falcon tube and the pelleted mitochondria were resuspended gently in 1 ml of SH buffer (0.6 M sorbitol, 20 mM HEPES, pH 7.5) and washed twice by centrifugation at 10,000 x g) for 15 minutes at 4°C to remove residual cytosolic contamination. Finally, the isolated mitochondria were resuspended in 50-100 𝜇l of SH buffer and quantified using Pierce BCA kit. The mitochondria were aliquoted and stored at -80°C until further use.

**Protein labelling for Import Assay**

To label the protein sample with a fluorescent dye for in-vitro import assay, Alexa Fluor™ C5 maleimide was used. The dye was added to the protein sample and incubated at 4°C for 3 hours in a dark place. To remove any unbound dye, NAP-5 columns were employed. Finally, the sample was concentrated using an amicon filter and stored at -80°C in multiple aliquots for future use.

**In-vitro Import Assay**

Firstly, the labelled protein should be unfolded (using 8 M Urea buffer) before the import assay is performed. 2 μl of labelled protein was mixed with 8 μl of 10 M Urea (resuspended in 1 X PBS) in an Eppendorf tube. The tube was covered with aluminium foil to protect the sample from light. The mixture was then incubated at room temperature for 1 hr. This unfolded protein was subsequently used for the mitochondrial import experiment. To set up the in-vitro import assay, the isolated mitochondria were diluted with 1X IA buffer (80 mM KCl, 5 mM MgCl2, 2 mM KH2PO4, 250 mM Sucrose, 10 mM MOPS-KOH, 5 mM DDT) at a ratio of 1:10. 25μg of isolated mitochondria was used for each reaction. To the sample, 1X PIC (protease inhibitor cocktail) and PMSF (Phenylmethylsulphonyl fluoride) were added at a final concentration of 2 mM. ATP and NADH were added to a 1:1 ratio at a final concentration of 10 mM. The mixture was incubated at 30°C for 3 minutes at 600 rpm. The unfolded protein (0.5μl) was introduced into the mixture, and the entire mixture was further incubated at 30°C for 30 minutes at 600 rpm. To stop the reaction, the tubes were immediately placed on ice. After centrifugation at 10,000 × g for 10 minutes at 4°C, the supernatant containing the proteins of interest was collected by adding 5X loading dye and boiling at 95°C for gel electrophoresis. The pellet was washed twice with 180μl of 1X IA buffer, followed by centrifugation at 10,000 × g for 10 minutes at 4°C. For gel electrophoresis, the pellet was resuspended in 1X loading dye and boiled at 95°C, similar to the supernatant. The gels were visualized on Typhoon FLA 7000 and band intensity were quantifed on ImageQuant TL.

**Probing the ubiquitination profile of the proteins**

For checking the ubiquitination profile of the proteins, the secondary cultures were inoculated at 0.2 O.D_600nm_ in 10ml of SD-ura from an overnight-grown primary culture, treated with AZC for 48 hours (chronic treatment). The cultures were allowed to grow till they reach O.D_600nm_ of 0.4-0.6. Once reached, the cultures were harvested and the protein was isolated using the alkaline lysis method. Protein estimation was done using Pierce^TM^ BCA kit (Thermo Scientific, Cat no-23225) and samples were prepared for western blotting. After transfer, the membrane was blocked with 2% BSA and decorated with Mono and Poly-Ubiquitin conjugated antibody (Enzo, BMLPW8810) for 16 hours at 4°C. The membranes was incubated with anti-rabbit antibody and were developed using Immobilon Western Chemiluminescent HRP substrate (Millipore, WBKLS0500) in the SYNGENE gel documentation system.

**Quantification And Statistical Analysis**

Student t test and R package for non-linear regression was used for statistical analysis. Mann-Whitney-U test was used for pathway specific analysis using Wilcox package in R.

**Data And Software Availability**

All data are provided in the manuscript. Genome sequencing files are under bioproject accession PRJNA1126056

**Key Resorce Table**

| **REAGENT** | **SOURCE** | **IDENTIFIER** |
| --- | --- | --- |
| **Antibodies** |  |  |
| Anti-NAT | (Ghosh et al., 2019) |  |
| Anti-Tim23 | Gift from Patrick D’silva Lab |  |
| Anti-mono and Poly Ubiquitin conjugate | Enzo | BMLPW8810 |
| Anti-Mouse secondary | Santa Cruz  Biotechnology | SC2005 |
| Anti-Rabbit secondary | Santa Cruz Biotechnology | SC2030 |
| **Bacterial Strains** |  |  |
| E.coli DH5𝛼  [F–, endA1 glnV44 thi-1 recA1 relA1 gyrA96 deoR nupG purB20 φ80dlacZΔM15 Δ(lacZYA-argF)U169, hsdR17(rK–mK+), λ–] |  |  |
| **Yeast Strains *( S. cerevisiae)*** |  |  |
| BY4741  (MATa his3Δ1 leu2Δ0 met15Δ0 ura3Δ0) | ATCC | ATCC#:201388 |
| W303 |  |  |
| RM11-1a |  |  |
| All yeast deletion strains used in this study | Invitrogen | 95401.H2 |
| All yeast DAmP strains used in this study | (Breslow *et al.*, 2008) | YSC5090 |
| All yeast GFP tagged strains used in this study | (Huh et al., 2003) | Invitrogen |
| **Oligonucletides** |  |  |
| mtCOX3_F_RT  TTGAAGCTGTACAACCTACC | (Rao et al., 2022) |  |
| mtCOX3_R_RT  CCTGCGATTAAGGCATGATG | (Rao et al., 2022) |  |
| nHsp31_F  ATGGCCCCAAAAAAAGTTTTACTCG | (Rao et al., 2022) |  |
| nHsp31_R  TCAGTTTTTTAAAGCGTCGATGGATC | (Rao et al., 2022) |  |
| nHsp31_RT_R CCGTTTCGTCTTGACCAT | (Rao et al., 2022) |  |
| **Chemicals** |  |  |
| NTC or ClonNAT | Jena Bioscience | AB102XL |
| AZC (L-Azetidine-2-carboxylic acid) | Sigma-aldrich | A0760 |
| G-418 disulfate salt | Sigma-aldrich | A1720 |

**Figure S1**

A) Growth rate of evolved strains at 30°C in YPD medium. TTB1, TTB2, TTB3, TTB4, TTB5, TTB6, TTB7, TTB8, TTB9, TTB10, TTB11 correspond to evolved thermotolerant strains and BY4741 is the parent strain.
B) Ratio of fluorescent (TDH2-GFP) and unfluorescent BY4741 population at 40°C at different time-points.
C) Histograms showing two distinct peaks corresponding to BY4741 and BY4741 (TDH2-GFP tagged) after mixing both cultures in a competitive fitness assay at 40°C for 24 hours (left panel). The right panel demonstrates the same for two different time points as indicated by colours.
D) Histograms showing the peak fluorescence in only BY4741 (left) and only GFP tagged TDH2 in BY4741 (right) at 40°C.

**Figure S2**

Graphs illustrating the complete loss of mtDNA in thermotolerant strains tby evaluating the depth of coverage at every locus of mitochondrial genome. Each graph rcorresponds to a specific strain, as indicated in the legend.

**Figure S3**

A) Relative fitness of evolved strains at 40°C after 100 generations of evolution. Fitness is normalized with the parent strain BY4741 . Mean is plotted with standard deviation as error bars from three biological replicates. Significance is calulated using Student’s t-test, p-value< 0.05.
B) Spot assay showing the growth benefit of evolved strains at 40°C as compared to BY4741.
C) Abundance of mitochondrial gene COX-3 in evoρ^0^ strains as compared to BY4741. Error bar represents standard deviation from three biological replicates. p-value<0.05
D) Oxygen consumption rate of a representative evoρ^0^ strain and the parent strain BY4741. Cells were added to the fresh mediumin the chamber and consumption of oxygen was monitored.
E) Spot assay of intermediate generations of TTB1 from 100 to 600 generations, showing the association of loss of mtDNA and thermotolerance. Cells were spotted on YP + 2% glycerol and YPD, incubated at 30°C and 40°C respectively.
F) Spot assay showing the impaired respiratory function in the evolved strains of W303 (W1, W2, W3) (left panel) but not in the evolved strains of RM11-1a (R1,R2, R3) (right panel).

**Figure S4**

A) Histogram depicting the increased expression of SSA4 and HSP12 upon treatment with AZC. SSA4 and HSP12 are fused with GFP and florescence of GFP is used as a proxy for expression of the chaperones. Gray arrow indicates shift in florescence of GFP upon treatment with 4mM AZC.
B) Median florescence of GFP is plotted where error bars depict standard deviation from three biological replicates. Significance is calculated using Student’s t-test, p-value<0.05.
C) Western blot showing increase in ubiquitinated proteins upon AZC treatment in BY4741. 30μg of total protein from cells treated with 4mM AZC or no AZC is loaded and probed with anti-mono and poly ubiquitin antibody. Replicates are indicated in the blot.
D) Confocal microscopy images illustration aggregation in BY4741 upon AZC treatment. Cells were treated with 4mM AZC for 6 hours and immobilized on chambered slides. GFP signal corresponds to HSP104 fused with GFP, a disaggregase of *S. cerevisiae* and a well-established marker of aggregation.
E) Relative fitness of TTB1 in 4mM AZC. Mean is plotted with standard deviation as error bars from three biological replicates. Fitness of TTB1 is normalized wrt BY4741. Significance is calculated using Student’s t-test, p-value< 0.05.
F) Abundance of mitochondrial gene COX-3 in intermediate generations (from 10 generation to 100 generation) of evolved strains. Error bars indicate standard deviation from three biological replicates. p-value<0.05
G) Spot assay showing the gain of AZC tolerance in the intermediate generations.
H) Spot assay showing the better growth of BYρ^0^ in AZC as compared to BYρ^+^. Log-phase culture were spotted on YPD, YP+ 2%glycerol (non-fermentative carbon source) and 5mM AZC, and incubated at 30°C for 48 hours.

**Figure S5**

A) Relative fitness of BYρ^+^ and BYρ^0^ strain in presence of mitochondrial translation inhibitors. C stands for Chloramphenicoland D stands for Doxycyclin. Two concentrations of each drug was used i.e., 35μg , 70μg of Chloramphenicol and 10μg, 20μg of Doxycyclin, along with 4mM AZC. Error bars indicate standard deviation from 3 biological replicates. p-value<0.05.
B) Growth assay of WT Nat-R and TS22 with varying concentrations of the antibiotic nourseothricin (Clonat) at 30 °C to assayfor the functionality of the proteins in intermediate generations of evoρ0 strains (20ρ^+^ , 20ρ^0^, 30ρ^0^ strain).
C) Growth assay of WT Nat-R and TS15 in BYρ^+^ and BYρ^0^ with increasing concentrations of nourseothricin (Clonat) at 30 °C.
D) The stability of WT Nat-R in BYρ^+^ and BYρ^0^, as determined by a cycloheximide chase at 30°C.
E) Cycloheximide chase assay of TS15 in BYρ^+^ and BYρ^0^ 30°C. Ponceau was used for normalisation of protein per lane (loading control) (n = 3).
F) Cycloheximide chase assay of TS22 in BYρ^+^ in chronic proteotoxic stress induced by AZC. Cells were treated with AZC for 48 hours and grown again in fresh medium till log-phase in presence of AZC.
G) Cycloheximide chase of ubiquitinated proteins in BYρ^+^ and BYρ^0^ to assess their degradation capacity at proteome-level. Figure shows the blot image (left) and the quatification of the same (right).

**Figure S6**

A) Mitochondrial fragmentation in BYρ^+^ upon treatment with 4mM AZC at 30°C (left panel). Quantification of the same is shown in right panel. Cells were transformed with plasmid containing mitochondrial-matrix targeted mCherry and were treated with 4mM AZC at 30°C. Images were collected after 6 hours.
B) MIC assay of BYρ^+^ and *∆doa1* in increasing concentration of AZC in YPD at 30˚C.
C) Relative fitness of *∆doa1* in 4mM AZC at 30°C, relative to BYρ^+^.
D) Cycloheximide chase assay of TS22 in BYρ^+^ (left panel) and BYρ^0^ (right panel) in the presence of 10μM Oligomycin. Ponceau is used for normalization of total protein.
E) in-vitro association of unfolded DMMBP with isolated mitochondria of BYρ^+^ and BYρ^0^. P corressponds to pellet fraction and S corressponds to supernatant fraction (left panel). Quantification of the same is shown in right panel.
F) In-vitro assay demonstrating the reduced degradation of unfolded DMMBP protein in isolated mitochondria of *∆tom70* comparedto BYρ^+^.
G) In-vitro assay demonstrating the reduced degradation of unfolded DMMBP protein in isolated mitochondria of BYρ^+^ in the presence of cytosolic domain of Tom70 CD and GroEL respectively.

**Figure S7**

A) Relative fitness of BYρ^0^ and BYρ^+^ in presence of single misfolded protein Fluc-DM. BYρ^0^ and BYρ^+^ cells expressing Fluc-DM were mixed with TDH2-GFP cells expressing Fluc-DM and outcompeting population was monitored by flow cytometry. Fitness of Fluc-DM is normalized wrt cells expressing WT Fluc.
B) PCR confirmation of the presence of COX-3 gene in *∆tom70* but not in *∆tom70*ρ^0^
C) Histogram depicting the expression of HSP104 in BYρ^+^ and BYρ^0^ and *∆tom70* upon treatment with 4mM AZC at 30°C. Fluorescence of GFP is used as a proxy for expression of HSP104 as HSP104 is fused with GFP.
D) Mitochondrial fragmentation in BYρ^0^ and *∆tom70* upon treatment with 4mM AZC at 30°C. Cells were transformed with plasmid containing mitochondrial-matrix targeted mCherry and were treated with 4mM AZC at 30°C. Images were collected after 6 hours.
E) Growth assay of TS22 in BYρ^+^ and *tim23* DAmP with increasing concentrations of nourseothricin (Clonat) at 30°C to assay for the activity of the protein. (n=3)
