## Supplementary figures and images for "Fitness defects due to cytosolic protein misfolding in *S. cerevisiae* can be alleviated by decreasing mitochondrial protein import capacity"

### Supplemental Figures

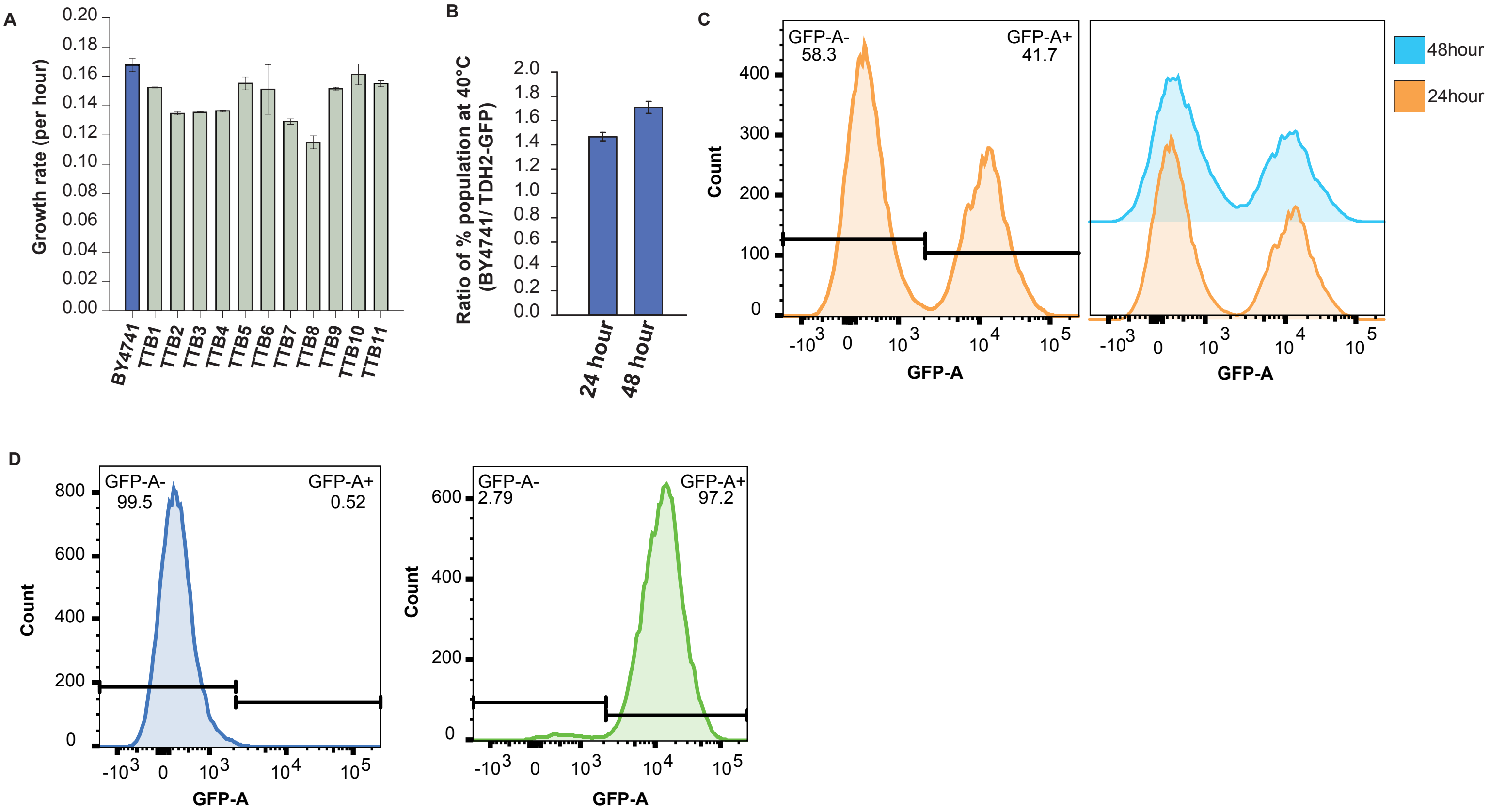

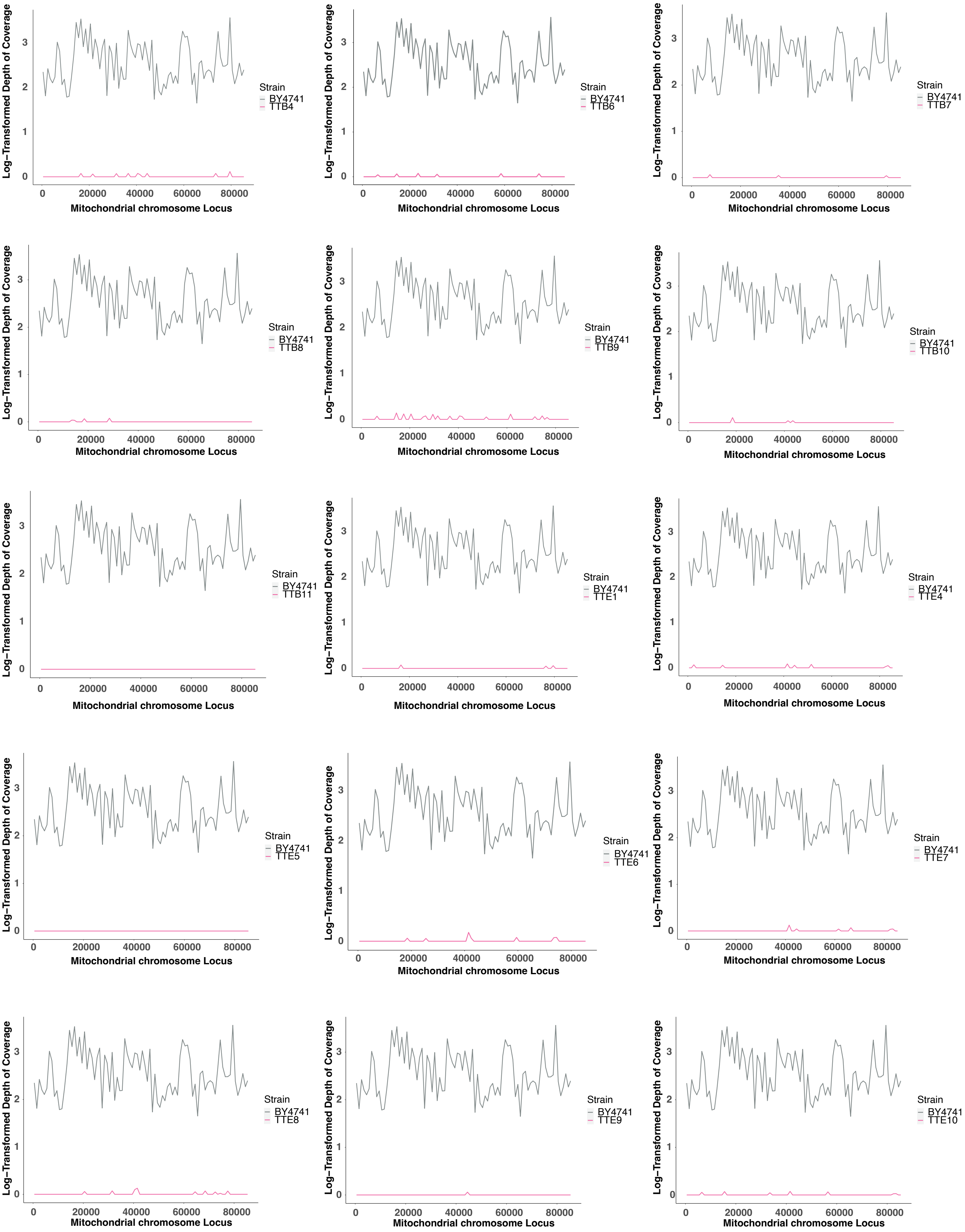

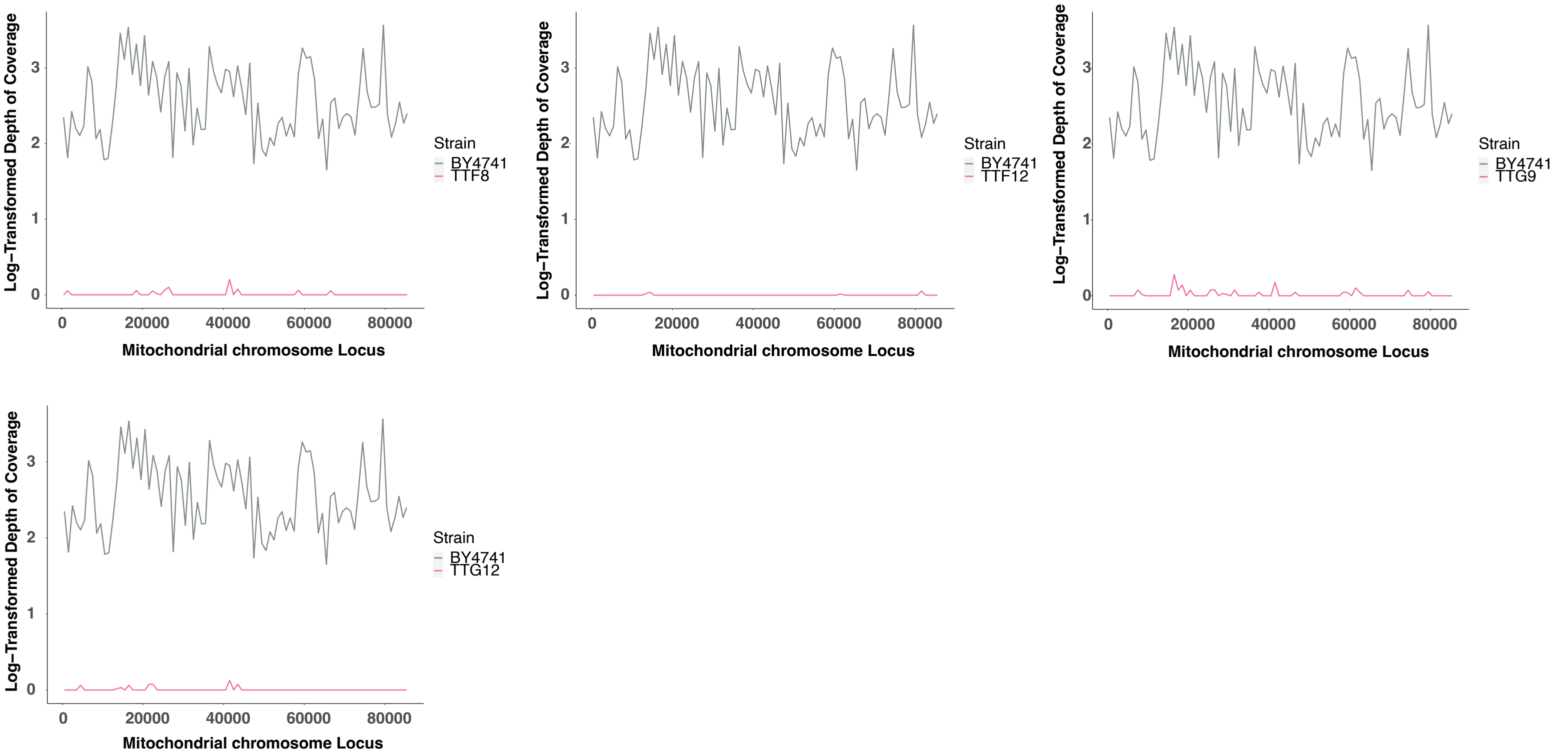

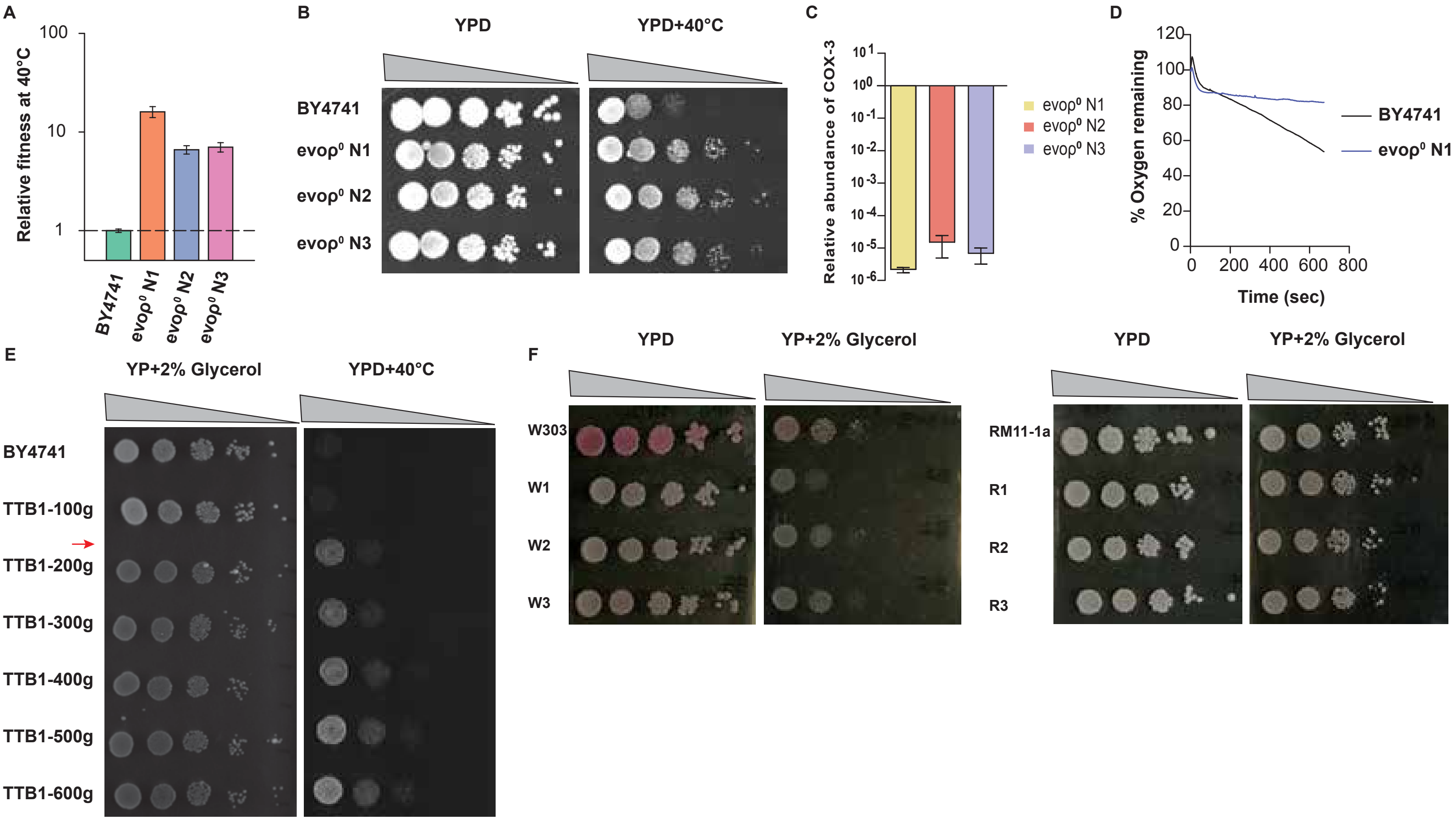

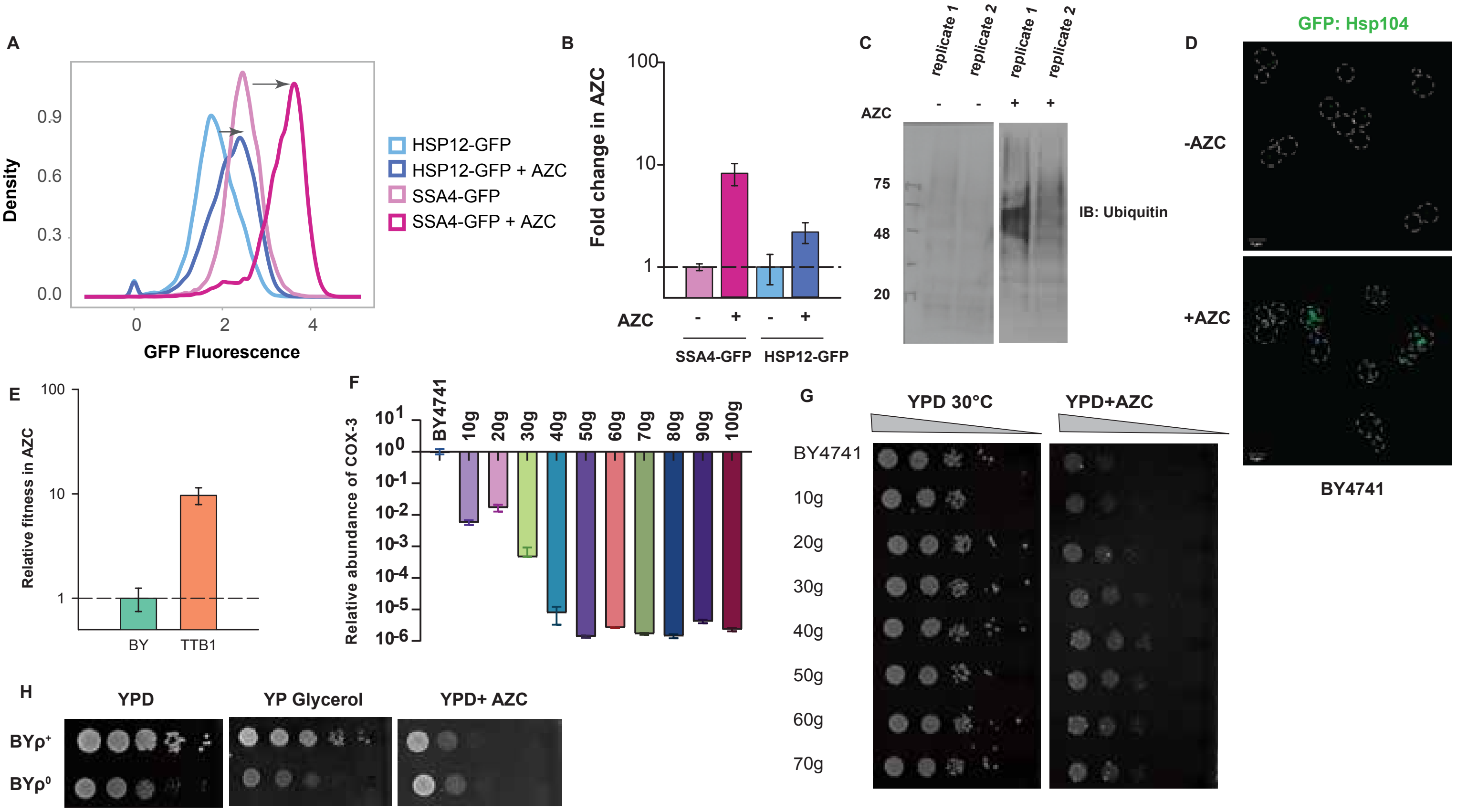

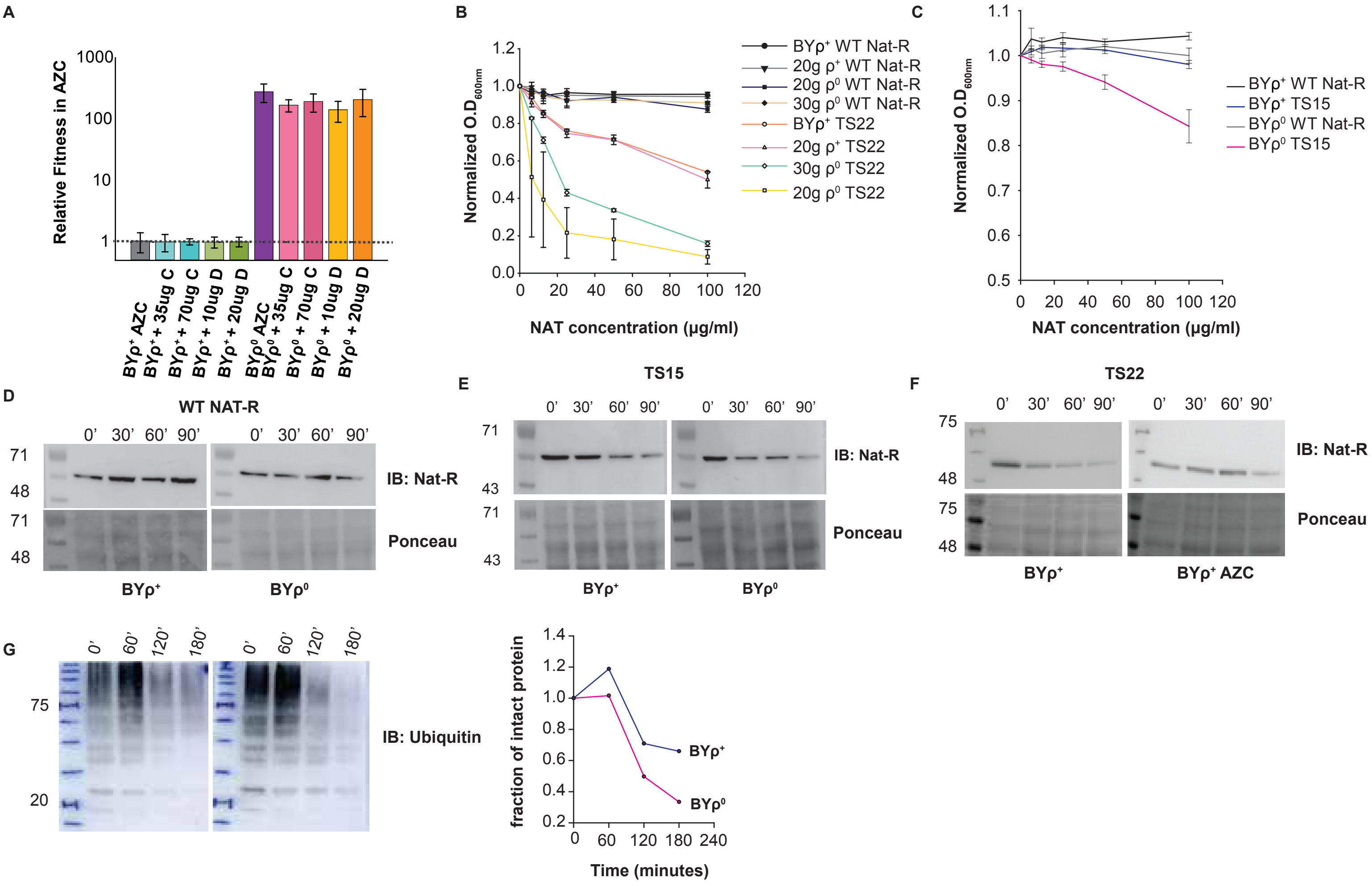

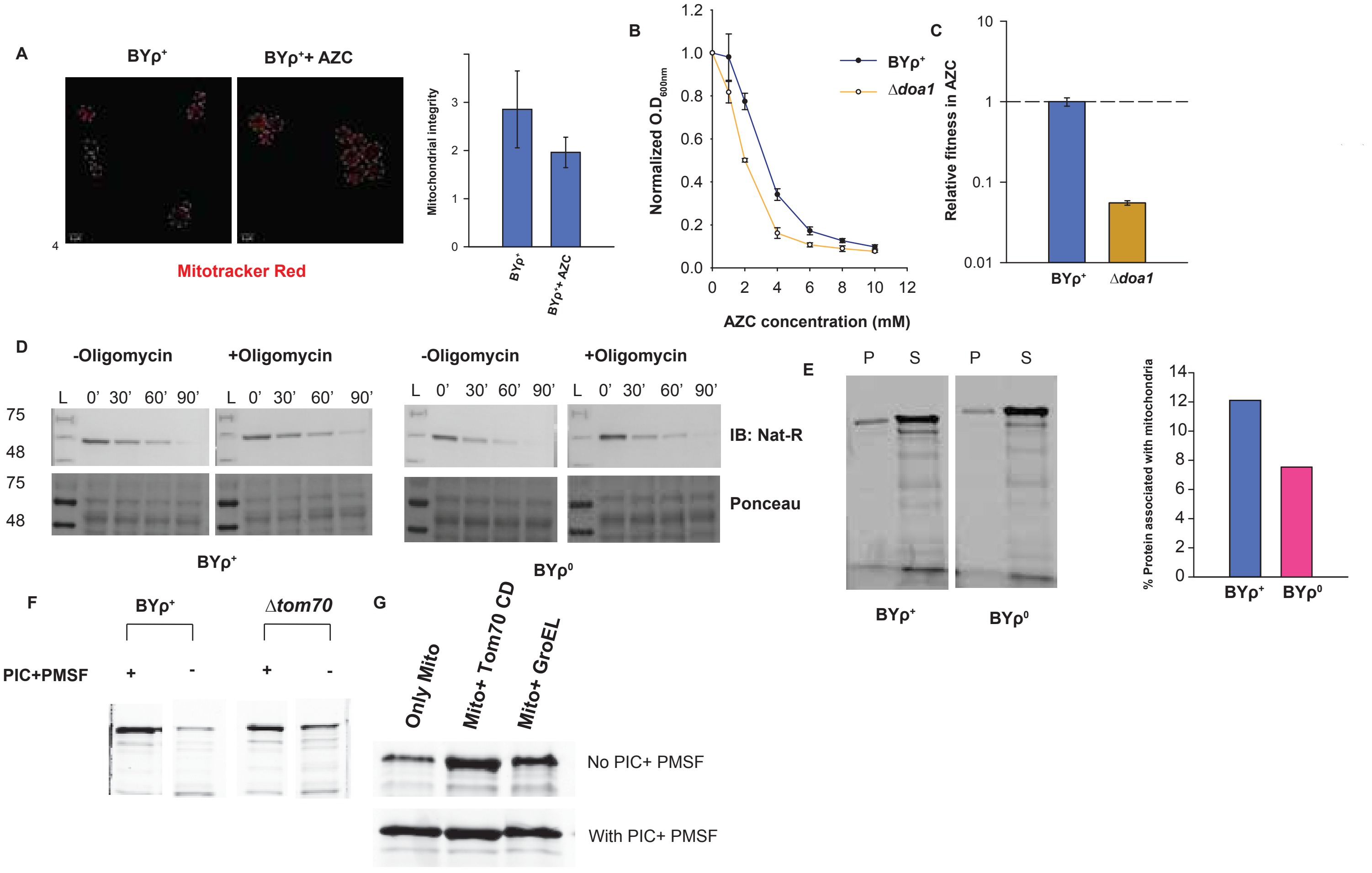

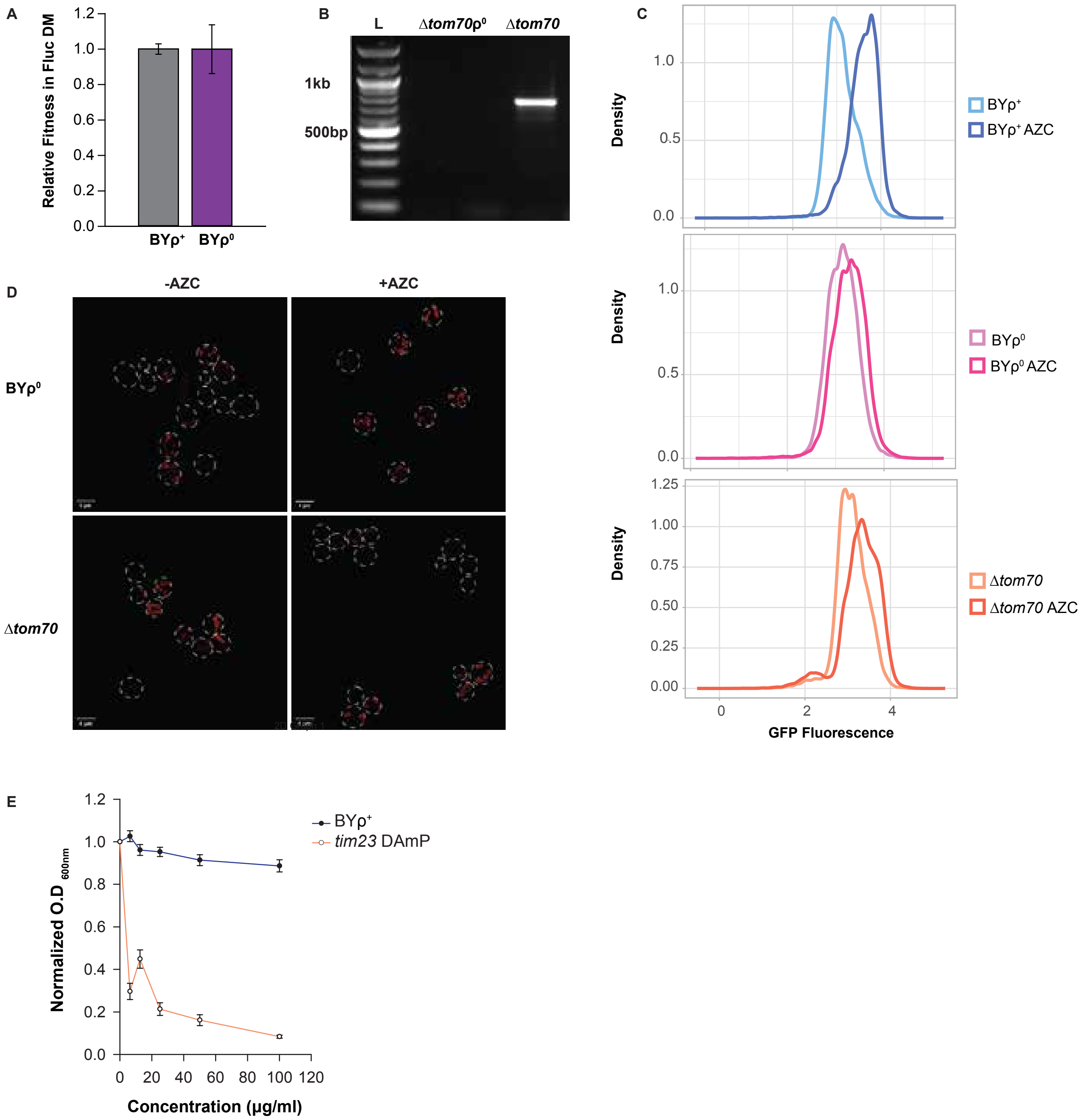
